# Rapid and robust directed differentiation of mouse epiblast stem cells into definitive endoderm and forebrain organoids

**DOI:** 10.1101/2021.12.07.471652

**Authors:** Daniel Medina-Cano, Emily K. Corrigan, Rachel A. Glenn, Mohammed T. Islam, Yuan Lin, Juliet Kim, Hyunwoo Cho, Thomas Vierbuchen

## Abstract

Directed differentiation of pluripotent stem cells (PSCs) is a powerful model system for deconstructing embryonic development. Although mice are the most advanced mammalian model system for genetic studies of embryonic development, state-of-the-art protocols for directed differentiation of mouse PSCs into defined lineages require additional steps and generate target cell types with lower purity than analogous protocols for human PSCs, limiting their application as models for mechanistic studies of development. Here, we examine the potential of mouse epiblast stem cells (EpiSCs) cultured in media containing Wnt pathway inhibitors as a starting point for directed differentiation. As a proof-of-concept, we focused our efforts on two specific cell/tissue types that have proven difficult to generate efficiently and reproducibly from mouse embryonic stem cells: definitive endoderm and neural organoids. We present new protocols for rapid generation of nearly pure definitive endoderm and forebrain-patterned neural organoids that model the development of prethalamic and hippocampal neurons. These differentiation models present new possibilities for combining mouse genetic tools with *in vitro* differentiation to characterize molecular and cellular mechanisms of embryonic development.

**SUMMARY STATEMENT:** New optimized protocols for directed differentiation of mouse epiblast stem cells into definitive endoderm and forebrain-patterned organoids.

## INTRODUCTION

Over the last two decades, methods for the directed differentiation of human PSCs (hPSCs) into numerous lineages have been developed and refined. As a result, it is now possible to generate nearly pure populations of multiple specific cell types or 3D multi-cellular organoids that resemble developing human tissues (Huch et al., 2017; Loh et al., 2016; Tchieu et al., 2017; Yiangou et al., 2018). However, while hPSC-based models for studying development advanced, analogous protocols for mouse PSCs have not yet emerged. Current state-of-the-art protocols for directed differentiation of mouse PSCs tend to be more complex than equivalent hPSC protocols, and they often fail to generate target cell types with high purity. Robust and efficient protocols for directed differentiation of mouse PSCs into defined lineages would be synergistic with *in vivo* studies in mouse and with studies of hPSC-based models of human embryonic development.

A critical difference between most mouse and human directed differentiation protocols is the starting cell state. hPSCs are generally cultured in the primed pluripotent state, which is roughly equivalent to cells of the pluripotent epiblast around the start of gastrulation (∼E6.5 in mouse) (Rossant and Tam, 2017). In contrast, mouse PSCs are predominantly grown in the naïve pluripotent state (mouse embryonic stem cells), which is equivalent to cells in the inner cell mass of the pre-implantation embryo at ∼E3.5-4.0 (Nichols and Smith, 2012). Like their embryonic equivalents, naïve PSCs must first exit the naïve pluripotent state before they can properly respond to differentiation cues (Morgani et al., 2017; Mulas et al., 2017; Smith, 2017). As a result, mouse naïve PSC directed differentiation protocols must include an additional first step of ∼48 hours in which cells exit the naïve pluripotent state and transition into an early formative and/or later primed pluripotent state that is competent to respond to signals that induce differentiation into specific somatic lineages. The requirement for this additional initial step is problematic because the naïve-to-primed transition is inherently asynchronous, leading to a heterogeneous starting population of formative/primed cells before directed differentiation has commenced (Kalkan et al., 2017; Strawbridge et al., 2020). The initial naïve-primed conversion step thus represents an impediment to the development of rapid and efficient protocols for directed differentiation that use mouse naïve PSCs as a starting population.

Mouse PSCs can be cultured in a primed pluripotent state, referred to as epiblast stem cells (EpiSCs). EpiSCs share many similarities with hPSCs, including the ability to respond rapidly to differentiation stimuli (Brons et al., 2007; Tesar et al., 2007; Vallier et al., 2009). Protocols that use EpiSCs as a starting point for directed differentiation have been developed, including (but not limited to) robust protocols for generation of oligodendrocyte progenitor cells (OPCs) and neuromesodermal progenitors (Edri et al., 2019; Najm et al., 2011). However, EpiSC-based protocols are not widely used for several reasons, including, 1) EpiSCs are prone to spontaneous differentiation and thus are more difficult to culture than mouse naïve PSCs or hPSCs, 2) EpiSCs have been reported to exhibit significant line-to-line variability in their differentiation propensity, and 3) EpiSCs express markers of lineage-primed epiblast cells transiting through the anterior primitive streak (T, FOXA2), which may limit their potential to differentiate into some lineages (Bernemann et al., 2011; Jouneau, 2019; Kojima et al., 2014; Kurek et al., 2015; Rossant and Tam, 2017; Song et al., 2016).

More recently, several groups demonstrated that inhibition of the canonical Wnt signaling pathway can limit spontaneous differentiation in EpiSC cultures, enabling consistent, long-term maintenance of pluripotent EpiSCs (Kurek et al., 2015; Sumi et al., 2013; Tsakiridis et al., 2014; Wu et al., 2015). Despite this progress, EpiSCs cultured in these conditions (+ Wnt inhibition; +WI) are still not commonly used for directed differentiation experiments.

Therefore, whether EpiSCs (+WI) have the potential to improve directed differentiation towards multiple lineages *in vitro* remains a critical gap in the field.

Here, we examine the potential of EpiSCs (+WI) as a starting point for *in vitro* directed differentiation studies. We focused on two specific cell populations that have proven difficult to generate reproducibly and efficiently from mouse naïve PSCs: definitive endoderm (DE) and neural organoids. First, we systematically optimized conditions to differentiate EpiSCs into DE, developing a protocol that converts EpiSCs into nearly pure DE in only 40 hours.

Second, we show that EpiSCs can robustly and reproducibly generate forebrain-patterned neural organoids. In depth characterization of forebrain-patterned organoids revealed that they undergo polarization into multiple distinct domains of progenitors, including prethalamic (P3) region of the caudal forebrain (diencephalon), hippocampus and cortical hem, as well as OPCs. The prethalamic progenitors, which gives rise primarily to inhibitory (GABAergic) neurons of the thalamic reticular nucleus and the zona incerta, are notable because they have not previously been efficiently generated in vitro (Shiraishi et al., 2017; Xiang et al., 2019). Our data suggest that mouse EpiSCs (+WI) can be a powerful platform for directed differentiation studies, enabling direct comparative studies with mouse embryonic development *in vivo* and with hPSC directed differentiation models.

## RESULTS

### Rationale for using EpiSCs (+WI) as a starting point for directed differentiation experiments

Given the issues associated with using mouse naïve PSCs for directed differentiation, we sought to assess the potential of mouse EpiSCs as a starting point for directed differentiation experiments. We chose to use mouse EpiSCs cultured with Wnt inhibitors (EpiSCs +WI), which had been previously shown to significantly reduce spontaneous differentiation and allow for maintenance of pluripotency over many passages in culture (Kurek et al., 2015; Sumi et al., 2013; Wu et al., 2015). These culture conditions have several features that make them potentially useful for directed differentiation experiments. First, EpiSCs (+WI) are easy for labs that currently work with mouse naïve PSCs to acquire, as naïve PSCs can be converted into stable EpiSC (+WI) lines in vitro. Second, there is suggestive data from previous studies indicating that EpiSC culture conditions work well across a variety of genetic backgrounds (including distinct inbred mouse strains) and with both male and female cells (Brons et al., 2007; Tesar et al., 2007). In contrast, culture of naïve PSCs from most inbred mouse strains requires addition of inhibitors of FGF/ERK signaling and GSK3 (2i culture conditions), which impedes *de novo* DNA methyltransferase activity, leading to erosion of parent-of-origin specific gene regulation (Choi et al., 2017; Czechanski et al., 2014; Yagi et al., 2017; Ying et al., 2008). Finally, given the similarities between EpiSCs (+WI) and hPSCs, we reasoned that it should be possible to adapt strategies developed for hPSC differentiation to EpiSCs (Vallier et al., 2009; Greber et al., 2010).

### Derivation and characterization of a panel of ground state EpiSCs

We derived EpiSCs (+WI) from 12 distinct naïve PSC lines, including lines from several distinct inbred mouse strains: C57Bl/6J (n=5), B6129SF1/J (n=3), DBA/2J (n=1), and PWK/PhJ (n=3) (Czechanski et al., 2014) (**Supplementary Table 4**). Briefly, naïve PSCs were converted into epiblast like-cells (EpiLCs) using previously established protocols (Hayashi et al., 2011). Following EpiLC conversion, cells were split onto irradiated mouse embryonic fibroblast feeders and maintained EpiSC media containing a Tankyrase inhibitor (NVP-TNKS656) to block canonical Wnt signaling (Shultz et al., 2013). Consistent with previous studies, EpiSC (+WI) express markers of post-implantation epiblast (OCT4, SOX2, OTX2, NANOG), do not express markers specifically expressed in naïve pluripotent stem cells (KLF4), and exhibit sparse expression of markers of primed epiblast (FOXA2) and primitive streak (T) (**Fig. 1A-B**).

**Figure 1:**
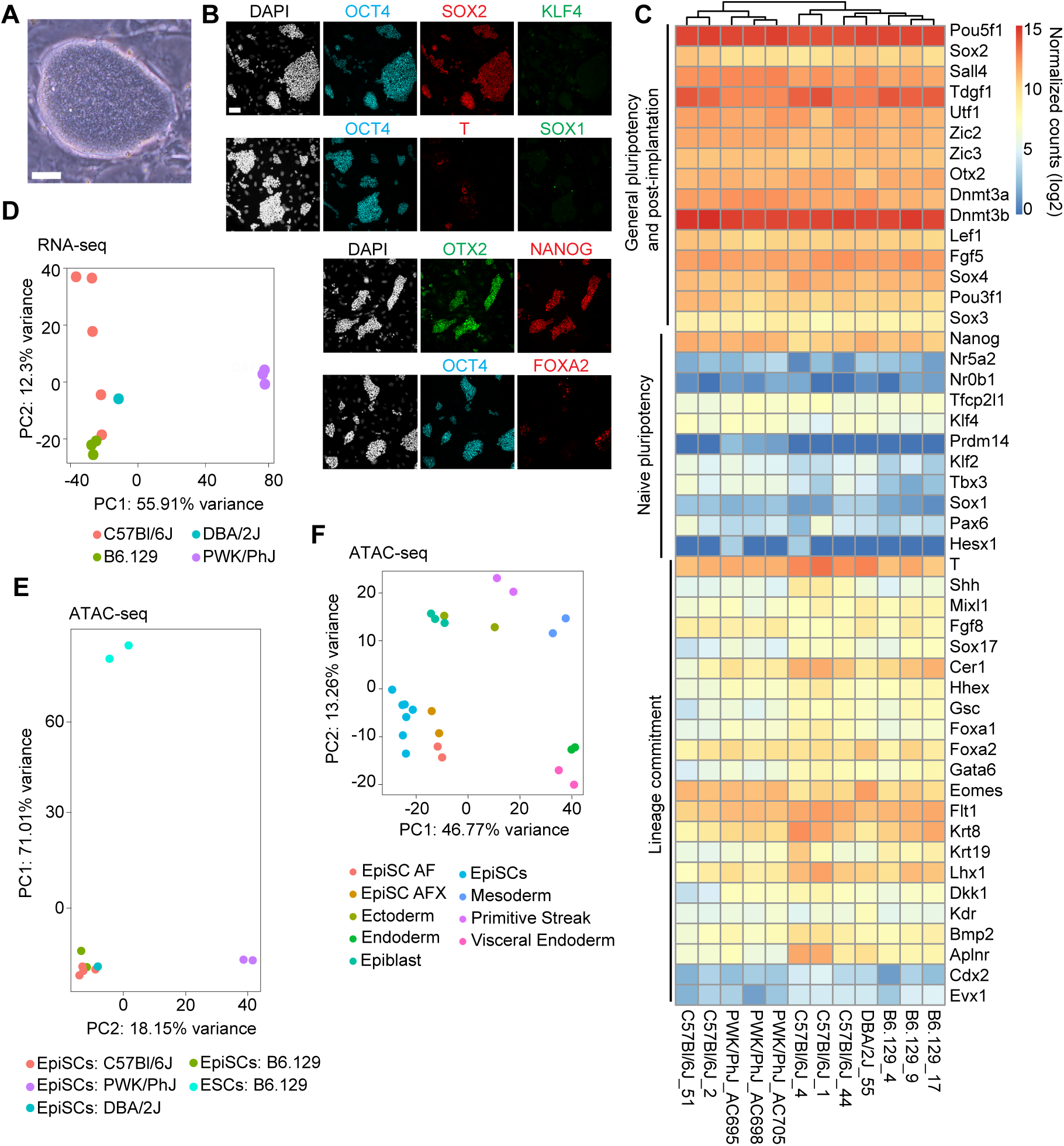
Derivation and characterization of mouse EpiSCs cultured in primed ground state conditions. (**A**) Phase contrast image of an EpiSC (+WI) colony. Scale bar = 50 μm. (**B**) Immunofluorescence images of EpiSC (+WI) cultures. Markers of general pluripotency (SOX2, OCT4), primed pluripotency (OTX2), naive pluripotency (KFL4, NANOG), and lineage commitment (T, FOXA2, SOX1). Scale bar = 50 μm. (**C**) Gene expression heatmap of RNA- sequencing data from EpiSC (+WI) lines (n=12; biological replicates derived from distinct naïve mESC lines) for selected genes associated with pluripotency and lineage-commitment. (**D**) Principal component analysis (PCA) plot of RNA-seq data (biological replicates; C57Bl/6J = 5, B6129SF1/J = 3, PWK/PhJ = 3, DBA/2J = 1) for EpiSC (+WI) lines. (**E**) ATAC-seq data (n =9 biological replicates; C57Bl/6J = 3, B6129SF1/J = 2, PWK/PhJ = 3, DBA/2J = 1) for EpiSC (+WI) and ESC lines (n=2; B6129SF1/J). (**F**) PCA plot of ATAC-seq data for EpiSC (+WI) lines (biological replicates; C57Bl/6J = 4, PWK/PhJ = 2, DBA/2J = 1). For comparison, ATAC-seq data from an additional set of EpiSC lines EpiSC-AF (cultured with Activin A and Fgf2) and EpiSC-AFX (cultured with Activin A, Fgf2, and Wnt inhibitor XAV939) from (Kinoshita et al., 2020) and from early embryonic lineages *in vivo* (E6.5 epiblast, E6.5 visceral endoderm, E7.5 ectoderm, E7.5 mesoderm, E7.5 endoderm, E7.5 primitive streak (from Xiang et al., 2020).

To characterize these EpiSC (+WI) lines in greater depth, we measured gene expression (RNA-seq; n =12 lines) and mapped accessible chromatin regions across the genome (ATAC-seq; n = 9 lines) (Buenrostro et al., 2015). We observed high expression of primed (*Otx2*, *Oct6, Dnmt3b)* and general (*Oct4/Pou5f1, Sox2, Nanog*) pluripotency markers, and low expression of naïve pluripotency markers (*Klf4, Tbx3*, *Prdm14*, *Nr5a2*) across all lines, in agreement with results from immunofluorescence staining (**Fig. 1B-C**). Similarly, genes associated with lineage commitment/lineage priming (*T*, *Foxa2*, *Mixl1*, *Sox17, Gata6, Cdx2, Sox1)* exhibited low to moderate expression across all lines (**Fig. 1C**). Principle component analysis (PCA) of RNA-seq and ATAC-seq data revealed that pluripotency state (naïve PSCs VS EpiSCs) is the primary driver of gene expression differences between lines, followed by genetic background (PC1, ∼56% and ∼71% of total variance, respectively, **Fig. 1D-E**). EpiSC lines from the same genetic background had highly correlated profiles of both gene expression and chromatin accessibility.

Next, we compared chromatin accessibility across the genome in our new EpiSC lines to cell populations in the embryo at embryonic day 6.5 (E6.5) (epiblast, visceral endoderm, primitive streak, and germ layer progenitors) as well as EpiSC lines cultured in the presence (EpiSC-AFX) or absence (EpiSC-AF) of the Tankyrase inhibitor XAV939 (Kinoshita et al., 2020; Xiang et al., 2020). PCA of chromatin accessibility profiles revealed that EpiSC lines most closely resemble EpiSC-AF and EpiSC-AFX, followed by *in vivo* epiblast tissue, and are least similar to primitive streak and endoderm/mesoderm) (**Fig. 1F**). Taken together, these data suggest that EpiSCs (+WI) exhibit expected features of primed PSCs and can be stably maintained in an undifferentiated state in culture.

### Optimized directed differentiation of EpiSCs into definitive endoderm

We next sought to test whether EpiSCs +(WI) can be directed to differentiate efficiently into DE. Current protocols for directed differentiation of hPSCs into DE are rapid and efficient, and serve as the foundation for robust protocols for generating progenitors of various endodermal organs (Yiangou et al., 2018). In contrast, protocols for DE differentiation of mouse PSCs require additional steps and give rise to heterogeneous populations of DE and mesodermal cell types (**Fig. S1A; Supplementary table 1**).

hPSC DE differentiation protocols consist of two steps that mimic the signals that sequentially specify DE during gastrulation (Gadue et al., 2006; Loh et al., 2014; Yiangou et al., 2018; Zorn and Wells, 2009). In the first step, Wnt and TGF-B/Nodal signaling promote exit from pluripotency and differentiation into anterior PS-like (aPS) progenitors. In the second step, high TGF-B/Nodal signaling promotes the commitment of aPS progenitors into DE. This general approach has been applied to a variety of hPSC lines and can consistently generate >85% pure DE (Loh et al., 2014). We hypothesized that signaling requirements for mouse DE differentiation from EpiSCs (+WI) would be similar to hPSCs, but that it was likely that the timing of each stage and the appropriate concentration of signaling factors would be different for mouse EpiSCs compared to hPSCs (Greber et al., 2010; Vallier et al., 2009)

To determine the optimal timing and concentration of each signal for aPS induction, we applied a gradient of increasing concentrations of CHIR99201 (to activate Wnt signaling via GSK3 inhibition) and Activin A (to activate TGF-B/nodal signaling) to EpiSCs (DBA/2J) for either 16, 20, or 24 hours (see methods for further details). After aPS induction, all conditions were shifted to the same DE commitment conditions for an additional 24 hours, and DE induction purity (fraction of the total population co-expressing FOXA2+/SOX17+) was measured using immunostaining (**Fig. 2A, S1B**). Across all conditions, we observed DE purity ranging from 7—82%. The highest purity was achieved in conditions with higher Activin A and the highest levels of Wnt activation. Exposure to these aPS conditions for 16 hours was sufficient to enable DE commitment in the second stage of the protocol, and no further increase in purity was observed in the 20 or 24 hour aPS conditions.

**Figure 2:**
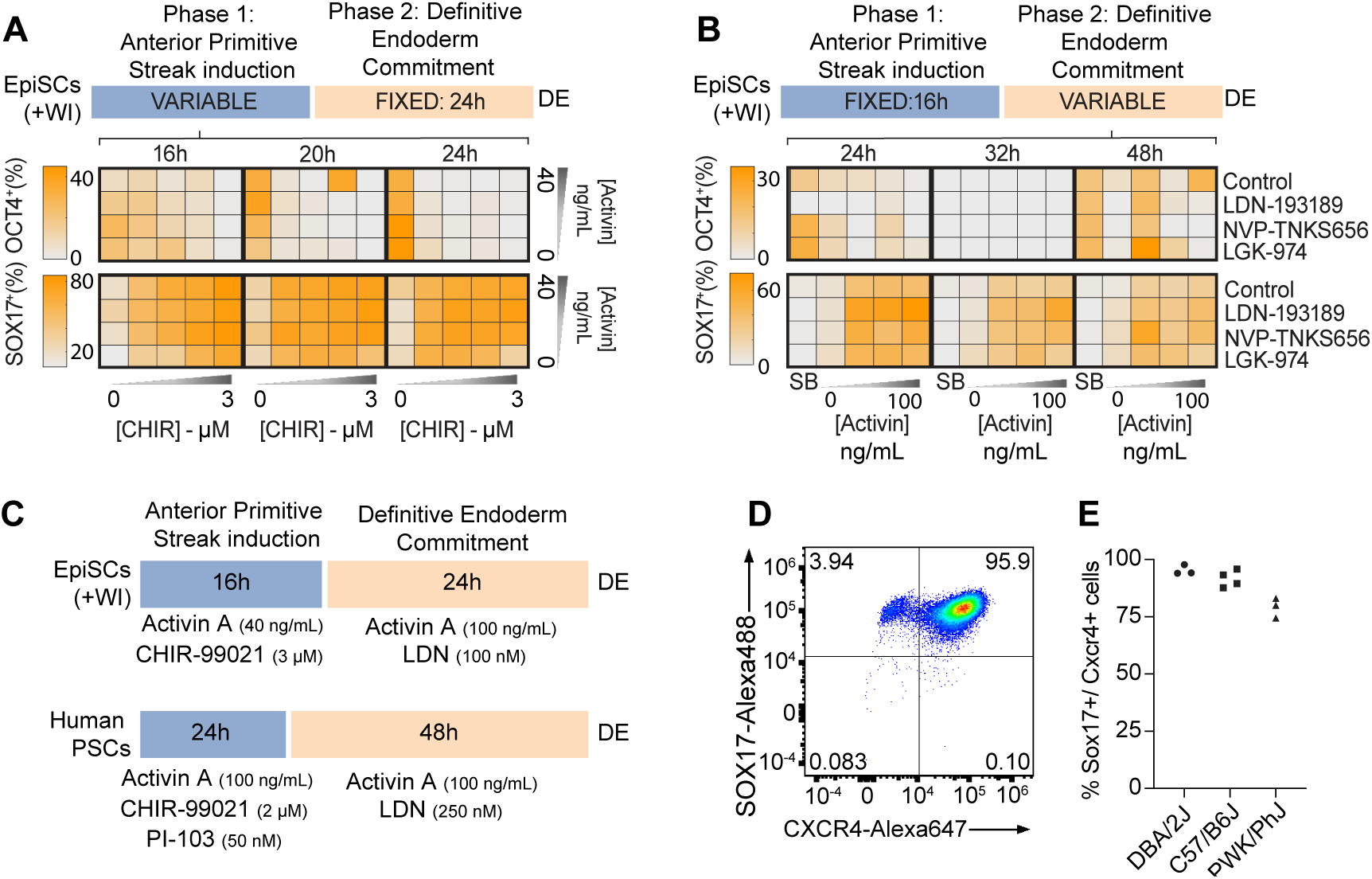
Systematic optimization of conditions for EpiSC-DE differentiation. (**A-B**) Results of DE protocol optimization experiments. For each condition, immunofluorescence staining was performed following 40 h of differentiation (end of Stage 2). Immunofluorescence images were quantified using CellProfiler. Markers of DE (FOXA2, SOX17), pluripotency (OCT4), and PS/mesoderm (T, CDX2) were examined. (**A**) Summary of results from anterior primitive streak (Stage 1) optimization (n=2 technical replicates). Note: We did not test the effects of < 16 hours of aPS induction. (**B**) Summary of results for optimization of DE commitment (Stage 2) (n=2 technical replicates). For Stage 1, the optimal conditions identified in (**A**) were used, then cells were switched into the indicated conditions for Stage 2. (**C**) Diagram of the optimized protocol for differentiation of mouse EpiSCs (+WI) into DE and a current hPSC to DE protocol (Loh et al, 2014). (**D**) FACS Quantification of immunostaining for DE markers (SOX17-Alexa488 and CXCR4-Alexa647) from cells at the endpoint of the optimized DE differentiation protocol. (**E**) Quantification of DE differentiation purity (SOX17+/CXCR4+ cells) across multiple experiments (technical replicates; C57Bl/6J, n=4; DBA/2J, n=3; PWK/PhJ, n=3).

Having defined these improved conditions for aPS induction, we next sought to define the optimal conditions for DE commitment of aPS-like cells (Stage 2) (**Fig. 2B, S1C**). To promote commitment of aPS-like cells into DE, DE differentiation protocols for hPSCs add a high concentration of Activin A for 48 hours. Concurrent inhibition of BMP and/or Wnt signaling has also been reported to reduce off-target differentiation of aPS cells into mesodermal lineages (Loh et al., 2014). Based on these data, we tested the effects of Activin A concentration, BMP inhibition (LDN-193189), Wnt inhibition (NVP-TNKS656, LGK-974), and the total duration of Stage 2 on DE commitment.

We induced aPS using the optimized conditions from **Fig. 2A** and examined the effects of different signaling conditions and timing (24, 36, 48 hours) on DE commitment. We observed the highest purity of DE differentiation in conditions with the highest levels of Activin A (∼75% of cells Sox17+ with 100 ng/mL vs. ∼65% Sox17+ cells with 40 ng/mL; Fig. 2B). Inhibition of BMP signaling markedly increased DE purity across all conditions, whereas Wnt inhibition had minimal effects. DE purity was highest in the 24 hour Stage 2 condition, with longer durations leading to a substantial reduction of FOXA2/SOX17 positive DE cells. In the 24 hour condition, rare SOX17- negative cells occur in small clusters of densely packed cells. Most of these off-target cells co-express OCT4 and T, are FOXA2 (-), and lack expression of CDX2, which is expressed by paraxial mesoderm cells that are also generated from aPS-like progenitors. These data suggest that the OCT4^+^/T^+^/FOXA2^-^ cells are most likely to be lagging aPS-like cells or immature mesodermal cells. However, in the 36 and 48 hour conditions, SOX17 (-) cells could also result from properly specified DE cells that have started to adopt distinct A-P identities that no longer express SOX17, such as anterior or posterior foregut (Li et al., 2018).

These systematic optimization experiments demonstrate that the combination of Wnt activation in the presence of moderate TGF-B/Nodal signaling, followed by TGF-B/Nodal activation and BMP inhibition can generate high purity DE from mouse EpiSCs in ∼40 hours (**Fig. 2C, Supplementary Experimental Methods 1**). Application of this protocol to EpiSCs (+WI) cultured in feeder-free conditions gave similar results (**Fig. S3B**).

To determine the purity of DE generated by our protocol more quantitatively, we used flow cytometry to measure the levels of the surface marker CXCR4 (expressed in endoderm and mesoderm) and the transcription factor SOX17 (expressed exclusively in DE) (**Fig. 2D**). This analysis indicated that the DE purity was ∼95% using the optimized conditions. Next, we used flow cytometry to assess the performance of our DE protocol on two additional EpiSC lines derived from distinct inbred strains of mice. The average DE purity achieved with a C57Bl/6J EpiSC line was similar to the DBA/2J line used for optimization (∼95%), whereas the purity achieved with a PWK/PhJ EpiSC line was slightly lower (∼80%) (**Fig. 2E**). However, for each line the results were similar between technical replicates from independent experiments (n=3-4 technical replicates/line).

### Molecular characterization of mouse EpiSC-derived definitive endoderm

To further characterize DE generated by our new protocol, we performed RNA-seq and ATAC-seq on DE generated from four independently derived C57Bl/6J EpiSC lines (characterized in **Fig. 1C-E**). CXCR4+ DE was purified with magnetic assisted cell sorting (MACS). EpiSC-derived DE exhibited large-scale transcriptional changes compared to EpiSCs, including upregulation of canonical DE marker genes (*Sox17*, *FoxA2*, *Cxcr4*, *Gata4/6*, *Cer1*) and down-regulation of pluripotency-associated genes, including *Oct4* (*Pou5f1*) and *Nanog* (**Fig. 3A, S2A-D**), consistent with results obtained using immunofluorescence staining (**Fig. S2E**) (Genga et al., 2019; Nowotschin et al., 2019). There are also widespread changes in chromatin accessibility in DE compared to EpiSCs. PCA of ATAC-seq data indicated that cell type (EpiSC vs. DE) accounted for the majority of variation observed across samples (PC1; 92.5%) (**Fig. S2B**). Accordingly, we observed significant increases in chromatin accessibility at numerous putative enhancer elements near genes known to be critical for DE differentiation (*Foxa2, Gsc, Otx2, Sox17*) (**Fig. 3B**). Across the genome, regions of hyper-accessible chromatin (peaks identified by ATAC-seq) in DE compared to EpiSCs are enriched for binding motifs for TFs known to play key roles in DE specification (GATA4/6, OTX2, SOX17, GSC). In contrast, ATAC-seq peaks in EpiSCs compared to DE exhibited enrichment for OCT4/POU5F1 and AP-1 (FOS/JUN) binding motifs (**Fig. 3C-E**). In summary, these genomic data demonstrate that DE exhibits global changes in gene expression and TF binding compared to EpiSCs and has the expected genomic features of *bona fide* PSC-derived DE.

**Figure 3:**
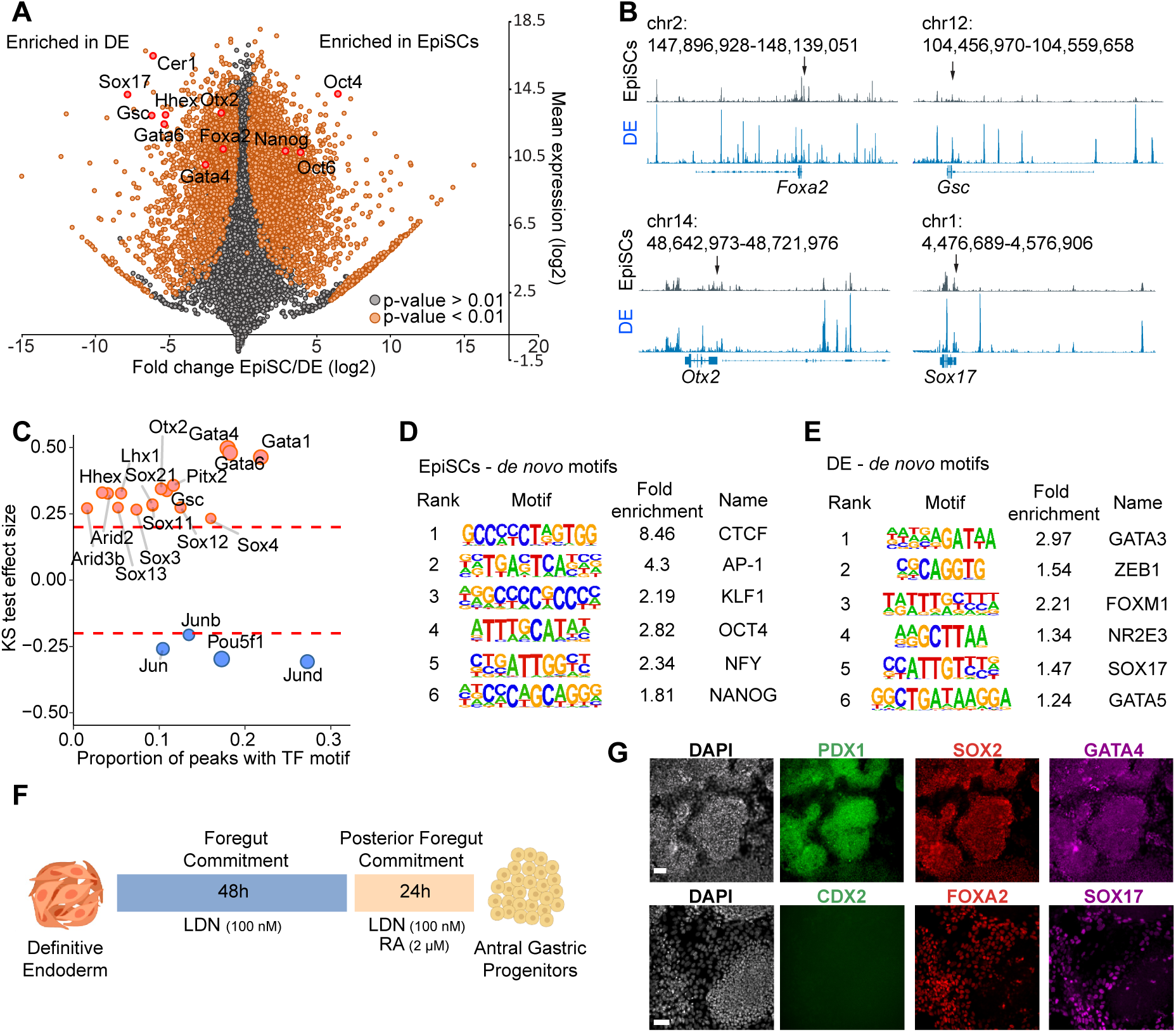
Genomic characterization definitive endoderm derived from EpiSCs (+WI). (**A**) RNA-sequencing data from C57Bl/6J EpiSCs (n=5, biological replicates) and C57Bl/6J DE purified via MACS (n=4, biological replicates). Differentially-expressed genes between EpiSC (+WI) and DE are highlighted in orange (P < 0.01, Wald test and corrected using the Benjamini-Hochberg method). A set of critical genes for primed pluripotency and DE are further annotated in red. (**B**) Chromatin accessibility (ATAC-seq) data from EpiSCs (+WI) (top row) or DE (bottom row). Gene loci known to play critical roles in DE development are displayed. Arrows indicate the promoter of each gene. (**C**) Comparison of TF binding motif enrichment in accessible chromatin regions during EpiSC (+WI) to DE transition. Peaks with differential chromatin accessibility between stages were compared to identify TF motifs enriched in cis-regulatory elements in each cell state (one-sided Kolmogorov-Smirnov (KS) test). TF motifs with a KS test effect size ≥ 0.20 are indicated by the dashed lines. (**D**) Results of *de novo* TF motif searches within EpiSC (+WI) or (**E**) MACS-sorted DE ATAC-seq peaks. (**F**) Diagram of the optimized protocol for differentiation of mouse DE into posterior foregut and antral gastric progenitors. Scale bar = 50 μm. (**G**) Antral gastric progenitors stained with antibodies against markers of posterior foregut (PDX1, SOX2 and GATA4), midgut/hindgut progenitors (CDX2) and definitive endoderm (FOXA2, SOX17) (n = 3 biological replicates). Scale bar = 50 μm.

Finally, we examined whether the DE generated by this protocol can be further patterned into specific populations of organ progenitors. We treated EpiSC-derived DE with conditions that promote posterior foregut patterning and subsequent differentiation into antral gastric progenitors (modified from protocols developed for hPSCs in (Broda et al., 2019; McCracken et al., 2014) (**Fig. 3F**). After 72 hours of additional differentiation, we observed a high density of cell clusters expressing markers of antral gastric epithelial progenitors (SOX2, PDX1, GATA4) and lacking expression of markers of midgut/hindgut (CDX2) and DE (FOXA2 and SOX17) (**Fig. 3G**). These data indicate that EpiSC-derived DE can be further patterned into organ progenitors and demonstrate the value of generating a high-purity population of DE for subsequent stages of differentiation.

### Efficient and homogeneous conversion of EpiSCs into forebrain-patterned organoids

The first neural organoid protocols were developed using embryoid bodies (EBs) formed from naïve mouse PSCs (Eiraku et al., 2008; Wataya et al., 2008). Given the success of our efforts to generate DE from EpiSCs (+WI), we next sought to determine whether EpiSCs (+WI) would also be a better starting cell type for generating neural organoids. We chose to focus our attempts on generating dorsal forebrain-patterned organoids, which have the potential to give rise to progenitors of multiple brain regions including the cerebral cortex, hippocampus, and thalamus (Montiel and Aboitiz, 2015).

Following extensive testing, we were able to identify conditions for robust EB formation, neural induction, and forebrain patterning of EpiSC-derived EBs (summarized in **Fig. 4A-B**). Briefly, EpiSCs (+WI) are dissociated into single cells and EBs are formed from ∼1,000 EpiSCs in neural induction media in Aggrewell plates. We found that addition of chroman 1 (ROCK inhibitor) and emricasan (pan-caspase inhibitor), components of the recently reported CEPT cocktail, help to dramatically reduce cell death during EB formation compared to commonly used ROCK inhibitors (Y-27632 or Thiazovivin; **Fig. S4A**) (Chen et al., 2021). To promote the generation of forebrain fates (telencephalon, diencephalon) we inhibited Wnt signaling via addition of the Porcupine inhibitor LGK-974 for the first two days of differentiation (Rifes et al., 2020; Tchieu et al., 2017). Inhibition of FGF signaling (via the FGFR inhibitor PD173074) during this period can promote forebrain patterning, but it is not essential for forebrain specification in this system. After the first 24 hours, EBs are removed from Aggrewell plates and embedded in Matrigel (based on Qian et al., 2018). Under these conditions, many EpiSCs express SOX1 after the first 24 hours of EB formation, and nearly all cells in EBs express SOX1 after 48 hours (**Fig. 4C**). In contrast, mouse neural organoid protocols that start from naïve PSCs have few SOX1+ cells after 72 hours in culture (Eiraku et al., 2008).

**Figure 4:**
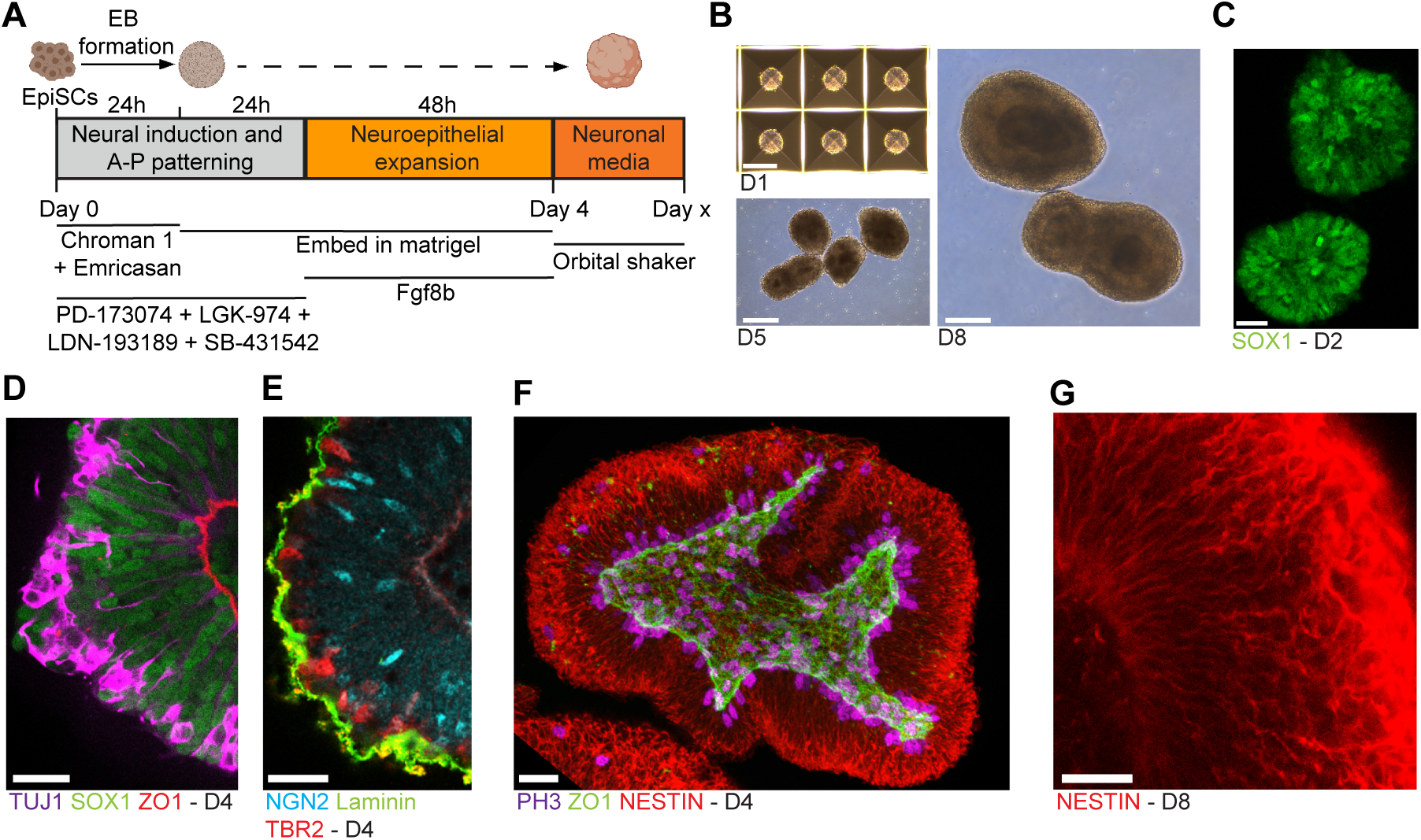
Generation of forebrain-patterned organoids from mouse EpiSCs. (**A**) Diagram of the protocol for forebrain-patterned neural organoid generation. (**B**) Brightfield images of forebrain organoids. Scale bar = 200 μm. (**C**) Confocal immunofluorescence image of developing organoid after 48 hours (d2). Scale bar = 25 μm. For **C-G**, all organoids were cleared (see materials and methods) and imaged *in toto* without sectioning. (**D-E**) Confocal immunofluorescence images of a d4 organoid. Scale bar = 25 μm. (**F**) Z-stack reconstruction of a d4 organoid. Scale bar = 25 μm. (**G**) Confocal immunofluorescence image of a day 8 organoid. Scale bar = 25 μm.

During early anterior-posterior patterning of the neural plate, FGF signaling promotes acquisition of more posterior neural fates, but shortly thereafter promotes telencephalic identity in the anterior-most region of the NE (∼E8.25) (Garel et al., 2003; Shimamura and Rubenstein, 1997). Therefore, we hypothesized that addition of FGF8 (FGF8b) after the first 48 hours of differentiation would facilitate induction of anterior telencephalic fates and could subsequently promote the proliferation and survival of telencephalic neuroepithelial progenitors (Storm et al., 2006). By 96 hours (d4), EBs form continuous neuroepithelia, with consistent apicobasal polarity (**Fig. 4D-E, S4B**) (**Video S1**). Characterization of the complete 3D structure of d4 organoids using tissue clearing revealed mitotic neuroepithelial progenitors at the apical membrane, a central lumen, Nestin-expressing cells with characteristic radial-glial morphologies spanning the length of the apical-basal axis of the developing tissue, as well as intermediate/basal progenitors and early-born postmitotic neurons migrating outward towards the basal lamina (**Fig. 4D-G**). Live-imaging of sparsely-labelled radial glial cells in organoids over a 24 hour period (d3-d4) revealed interkinetic nuclear migration and cell division behaviors characteristic of radial glial cells in the embryonic brain (Arai and Taverna, 2017; Noctor et al., 2004) (**Video S2)**. After d4, organoids are removed from Matrigel and switched into maintenance media (N2B27 media with vitamin A, and the neurotrophic factors BDNF and GDNF) in low adhesion cell culture dishes on an orbital shaker.

Next, we sought to characterize the specific regional identity of progenitors and neuronal populations generated in these organoids. At d4, all organoids expressed PAX6, which is expressed by progenitors in the caudal forebrain (diencephalon) and/or dorsal telencephalon, depending on the exact stage (Puelles et al., 2000). However, PAX6 expression was not homogeneous throughout the organoids, suggesting that not all neuroepithelial progenitors acquire the same regional identity under these conditions (**Fig. 5A**). By d8, most organoids spontaneously pattern into two opposing domains, which mirror the Pax6+ and Pax6- regions at d4 (Renner et al., 2017; Takata et al., 2017) (**Fig. 5B-G)**. Surprisingly, one domain contained large numbers of PAX6+ post-mitotic neurons, which occur almost exclusively in the developing prethalamus, generated from the anterior-most domain of the diencephalon (prosomere 3, p3) (Caballero et al., 2014; Manuel et al., 2015) (**Fig. 5C).** In the other domain, we observe neurons expressing markers of the cerebral cortex, such as TBR1, BCL11B/CTIP2, and BRN2, as well as RELN, a marker of Cajal-Retzius cells (**Fig. 5D-G, S5**). Taken together, these data suggest that organoids generated in these conditions polarize early into distinct domains that represent different anterior-posterior identities: the dorsal telencephalon and the anterior diencephalon (prosomere 3). Importantly, these results are reproducible across technical replicates and when starting with distinct EpiSC (+WI) lines (3 additional EpiSC lines) (**Fig. S5C-D, Supplementary Table 5**).

**Figure 5:**
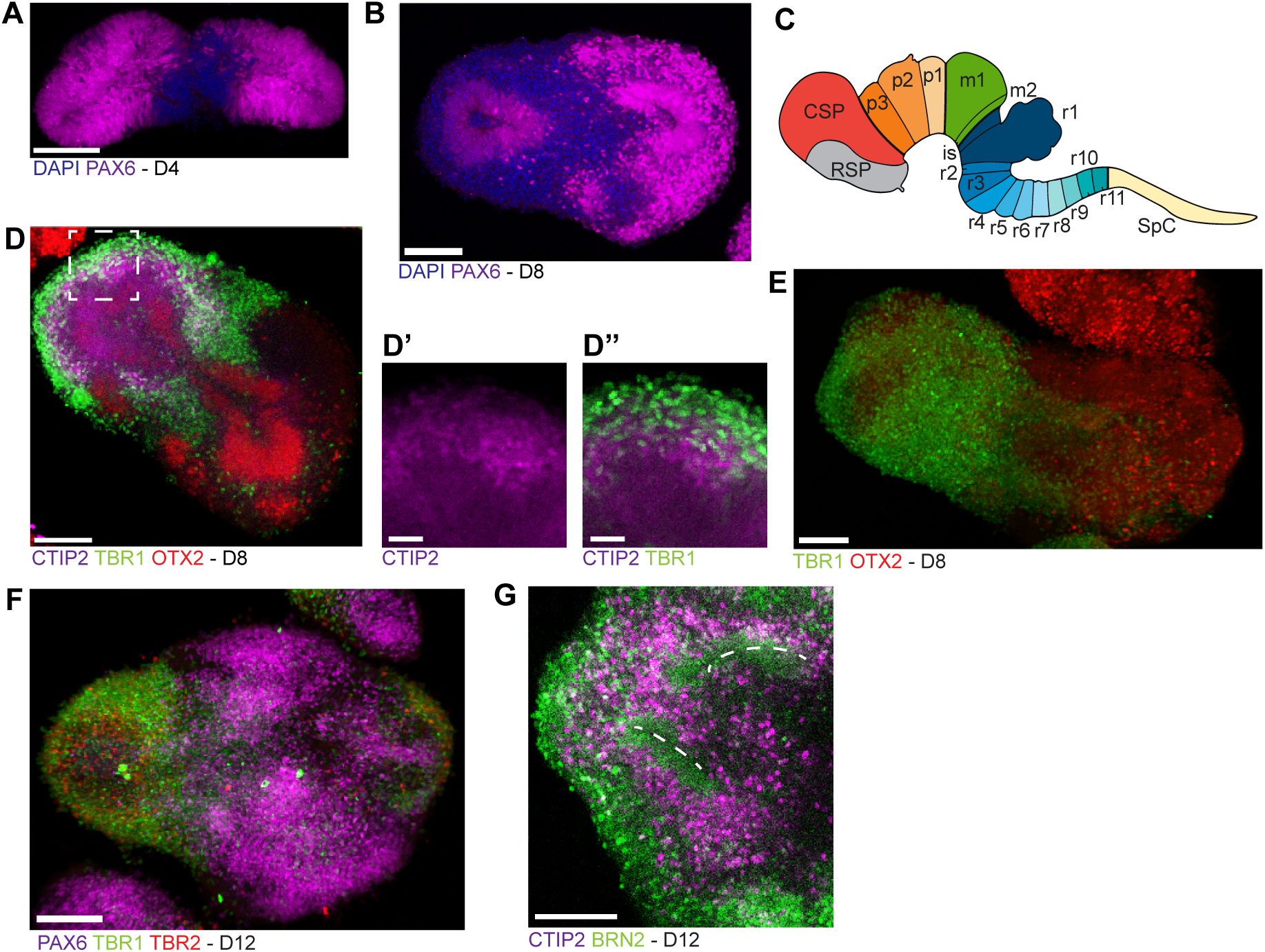
Characterization of the identity of progenitors and neurons generated in forebrain organoids. (**A-B**) Confocal immunofluorescence image of organoids at d4 (A) and d8 (B). Scale bar = 100 μm. (**C**) Diagram of prosomeres in the mouse brain (modified from Puelles et al., 2013). Abbreviations: rostral secondary prosencephalon (RSP), caudal secondary prosencephalon (CSP), prosomeres 1–3 of the diencephalon (p1, p2, and p3), midbrain mesomere 1 and 2 (m1 and m2), isthmic organizer (is), rhombomeres 1–11 (r1 to r11) and spinal cord (SpC). (**D**) Confocal immunofluorescence image of a d8 organoids (further magnified in D’ and D”), immunostained using antibodies against the deep layer cortical neuron markers TBR1 and CTIP2, and the forebrain neural progenitor marker OTX2. Scale bars (D) = 100 μm, D’, D” = 25 μm. (**E-G**) Confocal immunofluorescence image (3D projection) of a d8 (E) and d12 (F-G) organoids. Scale bar = 100 μm.

### Characterization of forebrain organoids using scRNA-seq

To gain further insight into the precise cell populations present within forebrain organoids, we generated scRNA-seq data from d12 organoids (5,652 total cells). Reduced-dimensionality visualization using UMAP revealed multiple distinct clusters, which we tentatively classified as specific cell types based on their expression of known marker genes. Importantly, nearly all cells within the organoid appear to be neuroectodermal derivatives, suggesting that neural induction in EBs is highly efficient (**Fig. 6, S6A, Supplementary table 6**)

**Figure 6:**
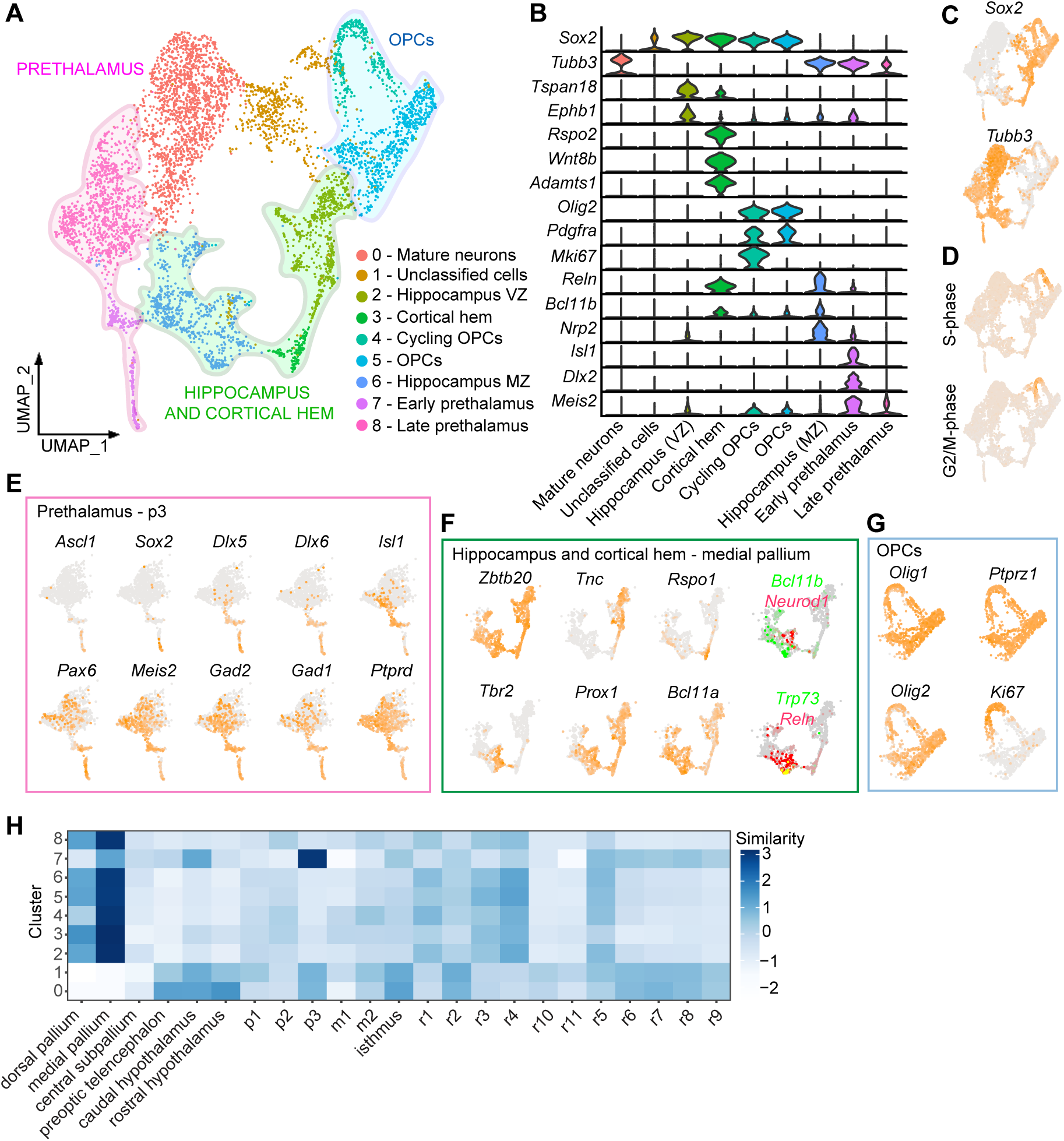
Characterization of day 12 forebrain organoids using scRNA-seq. (**A**) UMAP representation of scRNA- sequencing data from d12 organoids. Three broad classes of cell types are highlighted: oligodendrocyte progenitor cells (OPCs), hippocampus/cortical hem, and prethalamus. (**B**) Violin plots of gene expression levels for representative markers used to identify each cluster in (**A**) (see also **Supplementary table 5**). (**C**) Annotated UMAP highlighting the expression of *Sox2*, a neuronal progenitor marker, and *Tubb3* (Tuj1), a pan-neuronal marker. (**D**) Cell cycle analyses on d12 scRNA-seq data. (**E**) UMAP representations of cells from putative prethalamic clusters, highlighting expression of known pre-thalamic markers, from (**A**). (**F**) UMAP representations of gene expression data from putative hippocampal/cortical hem clusters from (**A**). Expression of *Ctip2* (*Bcl11b*) and *Neurod1*, which delineate the early development of the CA1-2 and CA3 regions of the developing hippocampus, respectively, appear to be mutually exclusive (Simon et al., 2012). (**G**) UMAP representations of gene expression data from putative OPC clusters from (**A**). (**H**) VoxHunt analyses on the scRNA-seq dataset, separated by cluster, compared to the E18 developing mouse brain. Abbreviations: prosomeres 1–3 of the diencephalon (p1, p2, and p3), midbrain mesomere 1 and 2 (m1 and m2), and rhombomeres 1–11 (r1 to r11).

Consistent with our immunostaining results, we observed clusters that appeared to be immature (cluster 7) and more mature (cluster 8) prethalamic neurons. In the embryo, the prethalamic domain generates multiple distinct types of GABAergic neurons that form the thalamic reticular nucleus and the zona incerta (Puelles et al., 2013). Accordingly, the early pre-thalamic neuron cluster (cluster 7) was characterized by expression of markers associated with early development of GABAergic neurons (*Ascl1*, *Dlx5*, *Dlx6, Gad1, Gad2*). The more mature prethalamic neuron cluster (Cluster 8) was characterized by expression of the prethalamic neuron markers (*Pax6*, *Isl1*, *Meis2*, *Mmp16)* (Mandai et al., 2014; Nagalski et al., 2016; Ono et al., 2014; Virolainen et al., 2012) (**Fig. 6E**). Within cells in this cluster, there was further heterogeneity based on the relative expression levels of *Ptprd* and *Pax6*, which distinguishes the lateral or the medial portions of the developing prethalamus *in vivo* (Guo and Li, 2019; Kim et al., 2020; Li et al., 2020; Sommer et al., 1997; Tuttle et al., 1999). In addition, more mature prethalamic neurons express higher levels of *L1cam* and *Nrcam*, which play a role in the pathfinding of axons from prethalamic neurons that project into the thalamus (Molnar et al., 2012; Quintana-Urzainqui et al., 2020) (**Fig. S6B**).

We identified one cluster (Cluster 6) that appeared to correspond to the glutamatergic neurons expressing TBR1 and/or BCL11B/CTIP2. However, although expression of TBR1/CTIP2 is often associated with cerebral cortex development, neurons in this cluster instead express several markers specific to hippocampal neurons (*Zbtb20*, *Neurod1*, *Prox1)*, and lack expression of the telencephalic marker *Foxg1*, which is expressed broadly in the cerebral cortex but absent from the hippocampus (Hatami et al., 2018) (**Fig. 6F)**. These data suggest that these neurons more closely resemble developing hippocampal neurons than deep layer cortical neurons. The observed similarity in marker profiles between these two cell types is consistent with *in vivo* data showing that the developing neocortex (cerebral cortex) and allocortex (hippocampus and olfactory bulb) have similar expression profiles during embryonic development (Cadwell et al., 2019).

We also observed a cluster of cells (Cluster 3) that resemble cortical hem, and are characterized by expression of *Rspo* genes, *Wnt8b*, *Wnt9a,* and *Bmp7* (Grove et al., 1998; Hasenpusch-Theil et al., 2012) (**Supplementary table 6**). The cortical hem is a secondary organizer located next to the developing hippocampus in the dorsomedial region of the developing telencephalon (Vieira et al., 2010). In addition to its role in cortical patterning via secretion of Wnt and BMP, neural progenitors in the cortical hem also generate Cajal-Retzius cells during early stages of telencephalon development (Takiguchi-Hayashi et al., 2004). Accordingly, we observe cells bridging clusters 3 and 6 that co-express the Cajal-Retzius cell markers *Trp73* (p73) and *Reln (reelin)* (Hanashima et al., 2007; Simon et al., 2012) (**Fig. 6F**).

We also used VoxHunt (Fleck et al., 2021) to comprehensively compare our single cell dataset to gene expression data from the developing embryonic brain (*in situ* hybridization data from the E18 Allen Developing Mouse Brain Atlas; Thompson et al., 2014). This analysis confirmed that the forebrain organoids most closely resemble medial pallium (hippocampus/cortical hem) and prethalamus/p3 (**Fig. 6H**).

In addition to these neuronal clusters, we also found clusters (Cluster 4 and 5) that appear to represent OPCs, characterized by high expression of *Olig1*, *Olig2*, *Pdgfra*, and *Dll1* (**Fig. 6G**, **Supplementary table 6**). Among all of the clusters, OPCs appear to be the only actively proliferating cells within organoids at d12 (**Fig. 6D**).

Immunofluorescence staining for OLIG2+ at d12 confirmed the presence of these cells, which were intermingled with CTIP2+ hippocampal neurons, suggesting that the same progenitors that give rise to the hippocampal glutamatergic neurons progress to produce OPCs (**Fig. S5C**). This is consistent with prior studies showing that mouse radial glial cells from the dorsal telencephalon give rise to OPCs after birth, and explant studies demonstrating that dorsal telencephalic progenitors from the embryonic brain often generate OPCs when explanted or cultured *in vitro* (Kessaris et al., 2006; Qian et al., 1997; Tekki-Kessaris et al., 2001).

Taken together, these data reveal that our EpiSC-based forebrain organoid protocol can robustly generate organoids containing multiple distinct regions of the developing forebrain, including dorsal, caudo-medial telencephalon (hippocampus and cortical hem) and dorsal anterior diencephalon (prosomere 3, prethalamus). These two domains are adjacent to one another, each bordering the boundary between the telencephalon and diencephalon. Prethalamic domains and hippocampal/cortical hem domains generally formed on opposite poles of organoids, suggesting that organoids undergo spontaneous polarization during early stages of develop and develop into distinct domains representing different anterior-posterior identities (Takata et al., 2017).

## DISCUSSION

Directed differentiation models enable a range of experimental techniques that would be difficult or impossible to perform *in vivo*, including genetic and chemical screens, live imaging, and experimental embryology approaches such as *in vitro* reconstitution (Hadjantonakis et al., 2020; Li et al., 2019; Moris et al., 2020; Schlissel and Li, 2020). Using mouse PSCs for directed differentiation has multiple advantages compared to hPSCs. For example, using mouse PSCs makes it possible to use the extensive genetic tools and resources that have been generated for mice. This includes not only knockout lines, reporter lines, and human disease models, but also numerous distinct inbred strains and outbred heterogeneous stocks such as Diversity Outbred mice that can be used for quantitative genetic approaches (Churchill et al., 2012; Skelly et al., 2020). In addition, results from mouse PSC differentiation models can be directly compared to genetically-matched tissues from mouse embryos, which allows for rigorous assessments of model validity (Voelkl et al., 2020). Such data represent an important proof-of-concept for hPSC-based cellular models of development and disease processes, as it will never be possible to perform similar cross validation experiments in developing human embryos (Kleiman and Engle, 2021).

Despite the advantages of mouse directed differentiation, mouse PSC models are much less frequently used than hPSC-based models. This is likely due in part to the limited availability of mouse PSC directed differentiation protocols that are robust, efficient, generalizable, and user-friendly. We found that by using EpiSCs (+WI), it is possible to develop protocols for directed differentiation as robust as state-of-the-art hPSC protocols. Since naïve PSCs can be converted into EpiSCs (+WI) *in vitro*, our new protocols can be applied to generate DE and neural organoids from pre-existing naïve PSC lines that carry specific genetic modifications, including gene knockouts or fluorescent reporters. These new optimized methods for directed differentiation represent a valuable addition to the experimental toolkit available for studying mouse development and thus should be of substantial interest to the developmental and stem cell biology research communities.

### Rapid and efficient generation of definitive endoderm from EpiSCs

By adapting protocols developed for hPSCs, we demonstrate that EpiSCs (+WI) can be differentiated into definitive endoderm with high purity (>90%) in only 40 hours. Our findings are consistent with previous work showing that EpiSCs and hPSCs respond similarly to most differentiation cues (Greber et al., 2010; Vallier et al., 2009). The most significant difference we observe between hPSC and EpiSC differentiation is timing. The optimal timing for EpiSCs is 16 hours for aPS induction and 24 hours to DE commitment, whereas hPSC protocols use 24 hours for aPS induction and 48 hours to DE commitment (Loh et al., 2014). These differences are consistent with the faster pace of mouse development compared to human, which has been previously observed when directly comparing mouse and human differentiation protocols (Ebisuya and Briscoe, 2018).

The high purity DE generated by our protocol will be an ideal starting point for subsequent differentiation into progenitors of a variety of endodermal organs, such as lung, liver, or intestinal tissues or organoids (Fowler et al., 2020; Yiangou et al., 2018). As a proof of concept, we demonstrated that EpiSC-derived DE can be further patterned to posterior foregut progenitors and antral gastric progenitors by adapting previously developed protocols for hPSC differentiation (Broda et al., 2019; McCracken et al., 2014). The high purity of DE achieved by this protocol should also facilitate large-scale genetic screens to identify genes that control endoderm commitment (Li et al., 2019).

### Improved methods for mouse neural organoid generation

The first neural organoid protocols were developed using naïve mouse PSCs (Eiraku et al., 2008; Wataya et al., 2008). Although these pioneering studies provided the foundation for the subsequent development of protocols to generate neural organoids from hPSCs, the original mouse protocols are now rarely used. This is due, at least in part, to the fact that these protocols were less robust and user-friendly than current hPSC organoid protocols. In addition, most of these early mouse neural organoid protocols pre-date the development of improved methods for consistently generating polarized neuroepithelia in neural organoids (e.g. matrigel embedding or addition of laminin/matrigel to the culture media) (Lancaster et al., 2013; Nasu et al., 2012). Therefore, we reasoned that by starting with EpiSCs instead of naïve PSCs, and by implementing improved techniques for organoid generation developed for hPSC, we would be able to develop an improved protocol for the generation of mouse neural organoids.

Although our initial goal was to generate a mouse organoid model of cerebral cortex development, we instead observed domains of hippocampal and prethalamic progenitors. This suggests that our conditions for anterior-posterior patterning must be further modified to generate cerebral cortex, which comes from the most anterior region of the developing neuroectoderm. Previous work with mouse neural organoids found that insulin exerts posteriorizing effects on developing neuroectoderm via activation of FGF signaling, and it is necessary to remove insulin from the media at early stages of neural induction to maintain anterior neuroectoderm patterning (Takata et al., 2017; Wataya et al., 2008). However, this strategy is not readily adaptable to primed PSC organoid models because insulin is critical for primed PSC growth and survival, making it difficult to form EBs and perform neural induction in insulin free conditions. Based on these studies, we added FGFR inhibitors in addition to Wnt inhibitors for the first two days to block the posteriorizing influence of FGF signaling. However, FGFR inhibition was not strictly required for forebrain specification if Wnt signaling is sufficiently inhibited. Within forebrain organoids, we observed spontaneous polarization into telencephalic and diencephalic regions (Renner et al., 2017; Takata et al., 2017). Interestingly, for hPSC-based cerebral cortex organoids, inhibition of Wnt signaling (in addition to dual SMAD inhibition for neural induction) is generally sufficient to maintain anterior neuroectodermal identity and specification of cortical fates, even in the presence of high levels of insulin. It will be interesting to explore the mechanisms underlying this apparent difference in anterior-posterior patterning between mouse and human.

Methods for the generation of organoids containing diencephalic progenitors have previously been reported (Shiraishi et al., 2017; Xiang et al., 2019). These protocols use a combination of moderate insulin, inhibition of FGF/ERK, and activation of BMP signaling (BMP7) to specify diencephalic progenitors of the thalamus and epithalamus (prosomere 2) and pretectum (prosomere 1). In contrast, we find that addition of high levels of FGF8b without exogenous BMP7 induces prethalamic (prosomere 3) fates. This is consistent with the high expression of FGF8b in the dorsal midline of p3 and prethalamic eminence at early stages of diencephalon development (E10.5- E12.5) (Martinez-Ferre and Martinez, 2009). *In vivo*, prethalamic progenitors generate numerous distinct types of inhibitory (GABAergic) neurons that form the thalamic reticular nucleus and the zona incerta (Li et al., 2020). Prethalamic neurons have not been generated efficiently in previous mouse and human thalamic organoids, and thus our forebrain organoid protocol should be useful for studying the development of this complex and understudied brain region.

Mouse forebrain-patterned organoids can complement *in vivo* mouse models of forebrain development as well as neurodevelopmental disorders (Sestan and State, 2018). Our neural organoid model should facilitate direct comparisons between the phenotypes observed *in vivo* in mice with mutations associated with neurodevelopmental disorders, and those observed in analogous mouse organoid models *in vitro*. In contrast, hPSC-based models of neurodevelopmental disorders cannot be conclusively validated *in vivo*, and thus it is difficult to evaluate the relevance of observations made in neural organoid models *in vitro* for developmental pathophysiology *in vivo*. More rigorous assessment of the strengths and limitations of *in vitro* models in mice can help to inform experimental design for hPSC-based modeling of neurodevelopmental disorders. In future studies, it will be interesting to perform in depth comparisons between developing neural organoids and their analogous cell types *in vivo*, and to compare mouse and human organoid development.

### Limitations of the study

Although we were able to develop robust protocols for differentiation of EpiSCs (+WI) into DE and neural organoids, it is possible that EpiSCs (+WI) will not be an ideal starting point for differentiation into all cell types. For some protocols, EpiLCs or more recently described stable formative stage stem cells might be superior to EpiSCs (+WI) (Hayashi et al., 2011; Kinoshita et al., 2020; Morgani et al., 2018; Yu et al., 2021). Even when cultured with Wnt inhibition, we observe some heterogeneity in EpiSC cultures, marked by expression of anterior primitive streak markers *Foxa2/T* (Bernemann et al., 2011; Blauwkamp et al., 2012; Song et al., 2016; Tsakiridis et al., 2014). Additional optimization of EpiSC (+WI) conditions could help to develop conditions that are as robust as hPSCs and further mitigate issues with current culture conditions such as residual heterogeneity and the need for fibroblast feeder cells. In this study, we exclusively used EpiSCs (+WI) that we derived in vitro via conversion of naïve mouse PSCs. Previous work suggests that EpiSCs derived in vitro are highly similar to EpiSC lines derived directly from post-implantation epiblast (Kojima et al., 2014), so we would expect cells from both sources to behave similarly, but we have not demonstrated that this is the case. There is an emerging consensus in the hPSC directed differentiation field that the initial culture conditions of hPSCs have a significant impact on their ability to differentiate into specific lineages (Cornacchia et al., 2019; Watanabe et al., 2019). Thus, it is likely that further improvements of EpiSC (+WI) culture will be the best way to generally improve their performance in directed differentiation experiments. The impact of long-term culture on genetic and epigenetic stability of EpiSCs (+WI) has not been characterized in depth (Halliwell et al., 2020). Similarly, the long-term effects of Tankyrase inhibition on the genetic and epigenetic stability of EpiSCs remain to be characterized. Several types of Wnt inhibitors have been used for EpiSC culture, and it remains uncertain how different means of inhibiting Wnt activity might change the properties of the resulting EpiSCs (Kurek et al., 2015; Sumi et al., 2013; Wu et al., 2015).

## MATERIALS AND METHODS

### Summary

Step-by-step protocols for directed differentiation experiments are provided as **Supplementary Experimental Methods 1**

For information about antibodies, see **Supplementary Table 2**

For reagent information, see **Supplementary Table 3**.

Detailed information for each cell line is provided in **Supplementary Table 4.**

Detailed information for each cell line used for forebrain organoid generation is provided in **Supplementary Table 5.**

### Complete UMAPs, related to figure 6, are provided in Supplementary Table 6. Mouse ESC culture and conversion to EpiSCs

All mouse ESC lines were maintained on gelatin-coated dishes with irradiated mouse embryonic fibroblast feeder cells using serum/LIF media comprised of DMEM (high glucose, GlutaMAX, HEPES), 1% nonessential amino acids, 1% sodium pyruvate, 1% penicillin-streptomycin, 0.1% 2-mercaptoethanol, 10% fetal bovine serum and 1000 units/mL ESGRO LIF. All ESC lines (except lines from B6129SF1/J background) were cultured in serum/LIF media with 2i containing 3 µM CHIR99201 and 1 µM PD0325901. Media was changed daily, and ESCs were passaged upon 70% confluence at a 1:6 ratio using TrypLE. ESC-to-EpiLC conversion was performed as described in (Morgani et al., 2018). Briefly, ESCs were lifted using 0.5 u/µL Collagenase IV, centrifuged and dissociated into a single cell solution using Accutase. The ESC suspension was plated on fibronectin coated plates (16.7 µg/mL) at a density of 17,500 cells/cm^2^ in N2B27 media supplemented with 12.5 ng/ml heat stable recombinant human bFGF, 20 ng/ml Activin A, and 1% Knockout Serum Replacement. N2B27 media consists of 50% DMEM-F12, 50% Neurobasal, 0.5% N2 supplement, 1% B27 supplement without vitamin A, 2 mM glutamax, 1% penicillin-streptomycin, and 0.1% 2-mercaptoethanol. The media was changed after 24 hours, and cells were converted to EpiSCs after 48 hours. For EpiLC-to-EpiSC conversion, EpiLCs were dissociated into small clumps (∼3-5 cells) with Accutase and plated at a density of ∼50,000-100,000 cells per cm^2^ on irradiated mouse fibroblast feeder cells in N2B27 supplemented with 20 ng/ml Activin A, 12.5 ng/ml heat stable bFGF and Wnt inhibitor (175 nM NVP- TNKS656). EpiSC media was changed daily, and cells were passaged every ∼48 hours at a 1:6 ratio using 0.5 u/ul collagenase IV followed by dissociation with Accutase into small clumps of 3-5 cells. For feeder-free EpiSC culture, EpiSCs growing on feeders were isolated using 0.5 u/ul collagenase IV and seeded on fibronectin-coated plates (16.7 µg/mL). After the first feeder-free passage, cells were passaged using Accutase only. The EpiSCs were considered as feeder-free after 2 passages with no visible feeder cell remaining.

### Definitive endoderm and antral gastric progenitor differentiation

For definitive endoderm differentiation, EpiSCs were generally used after ∼4-8 passages in EpiSC conditions. For differentiations, high quality cultures of EpiSCs were detached using Collagenase IV (0.5 u/μL), washed once with PBS, and then dissociated into a single cell suspension using Accutase. EpiSCs were plated at a density of 110,000 cells/cm^2^ in chemically defined media (CDM) containing 50% IMDM, 50% Ham’s F12 Nutrient Mix with Glutamax, 1% chemically defined lipid concentrate, 450 µM Monothioglycerol, 1% polyvinyl alcohol (w/v), 15 µg/ml Apo-transferrin, 0.5% Glutamax, 0.7 µg/ml Insulin and supplemented with 20 ng/ml Activin A, 12.5 ng/ml heat stable bFGF, 175 nM NVP-TNKS656, 1% knockout serum replacement, and 2 µM Thiazovivin. Plating media was removed 6 hours after seeding, cells were washed gently with PBS without calcium or magnesium (PBS-/-) and the media was changed to CDM supplemented with 3 µM CHIR99201 and 40 ng/ml Activin A. 16 hours after the first media change, cells were washed with PBS-/- and the media was replaced with CDM supplemented with 100 ng/ml Activin A and 100 nM LDN-193189. After 24 hours (46 total hours after cells were initially seeded), cells were fixed for immunostaining or collected for further analysis. For antral gastric progenitor differentiation, definitive endoderm cells were kept in CDM with 100nM LDN and 2% fetal bovine serum, with a renewal of the media after 24 hours. After 48h, the media was changed to CDM with 100nM LDN, 2 µM retinoic acid and 2% fetal bovine serum for extra 24 hours. After 72 hours cells were fixed for immunostaining.

### Immunostaining

For immunostaining of adherent cells (EpiSCs and DE), cells were rinsed twice with PBS-/- and fixed with 4% paraformaldehyde solution for 30 min at room temperature. After fixing, cells were washed twice with PBS -/- and permeabilized with PBS -/- containing 1% Triton-X (PBST) for 10 minutes at room temperature and blocked for 30 minutes with PBS -/- containing 0.3% Triton-X and 5% fetal bovine serum. Primary antibodies were diluted in PBST with 1% fetal bovine serum and incubated overnight at 4C. Detailed information about the antibodies used can be found in **Supplementary Table 2**. Following overnight incubation and three 10 min washes with PBST, cells were incubated with secondary antibodies for 2 h at room temperature. Cell were rinsed with two 10 min washes of PBST before DAPI staining (1 µM) for 15 min at room temperature. Cells were rinsed with PBST and imaged using a Leica Dmi8.

### Quantification of Immunofluorescence images

All image quantification was performed using Cell Profiler version 4.2.1 to calculate efficiency of DE differentiations via FOXA2 and SOX17 expression. DAPI-stained nuclei were identified using the Identify Objects module. To identify nuclei stained with each antibody, segmented nuclei were further analyzed using the Identify Objects module and the Relate Objects Module. These data were then classified as either positive or negative for each marker using the Filter Objects module and overall efficiency was calculated by percentage of expression for each marker.

### Flow Cytometry

Following DE differentiation, cells were dissociated in Accutase and washed in FACS buffer consisting of PBS-/-, 20% fetal bovine serum and 0.1% 0.5 M EDTA. CXCR4 antibody conjugated to Alex647 and live/dead zombie UV staining were diluted in FACS buffer at the manufacturer’s recommended concentration and incubated for 15 minutes at room temperature. Cells were washed and resuspended in FACS buffer with fixation/permeabilization diluent and fixation/permeabilization concentrate for 30 minutes at room temperature. The cells were then resuspended in permeabilization buffer with SOX17 antibody conjugated to Alexa488 at the manufacturer’s recommended concentration for additional 30 minutes at room temperature. Fixed cells were then analyzed using the Cytek Aurora.

Flow cytometry results were analyzed using FlowJo software v10.8 and single color stainings were used as a negative control for gating purposes.

### MACS purification of Definitive Endoderm

Definitive endoderm cells were isolated for ATAC-seq and RNA-seq analyses using MACS following manufacturer’s instructions, with slight modifications. Cells were dissociated in Accutase, washed in FACS buffer, counted, and resuspended with Rat anti-mouse CD184/CXCR4 Alexa Fluor 647 conjugated antibody at a concentration of 1 uL/10^6^ cells and incubated for 1 h at 4°C. Cells were then washed with FACS buffer, centrifuged, and resuspended in a solution composed by 80% FACS buffer and 20% Anti-Rat IgG MicroBeads. Cells were incubated in the mixture for 15 min at 4°C, then washed and resuspended in 500 uL buffer. The MS column was placed in OctoMACS separator on the MACS multistand. The cell suspension was added to the column and rinsed 3 times with buffer. Finally, the column was removed from separator and the magnetically labeled cells were collected.

### RNA extraction for RNA-seq

Phase separation in cells lysed in TRIzol was induced with 50% isopropanol + 0.5% 2-Mercaptoethanol and RNA was extracted from the aqueous phase using the MagMAX mirVana Total RNA Isolation Kit on the KingFisher Flex Magnetic Particle Processor according to the manufacturer’s protocol with 1-2M cells input. Samples were eluted in 38 µL elution buffer.

### Transcriptome sequencing of EpiSC samples

After RiboGreen quantification and quality control by Agilent BioAnalyzer, 100-500 ng of total RNA with RIN values of 9.5-10 underwent polyA selection and TruSeq library preparation according to instructions provided by Illumina (TruSeq Stranded mRNA LT Kit), with 8 cycles of PCR. Samples were barcoded and run on a NovaSeq 6000 in a PE100 run, using the NovaSeq 6000 S4 Reagent Kit (200 Cycles) (Illumina). An average of 36 million paired reads were generated per sample and the percent of mRNA bases per sample ranged from 84% to 89%.

### Transcriptome sequencing of Definitive Endoderm samples

After RiboGreen quantification and quality control by Agilent BioAnalyzer, 2 ng total RNA with RNA integrity numbers ranging from 7.7 to 10 underwent amplification using the SMART-Seq v4 Ultra Low Input RNA Kit, with 12 cycles of amplification. Subsequently, 10 ng of amplified cDNA was used to prepare libraries with the KAPA Hyper Prep Kit using 8 cycles of PCR. Samples were barcoded and run on a NovaSeq 6000 in a PE100 run, using the NovaSeq 6000 S4 Reagent Kit (200 Cycles) (Illumina). An average of 39 million paired reads were generated per sample and the percent of mRNA bases per sample ranged from 83% to 87%.

### ATAC-seq

ATAC-seq was performed as previously described (Buenrostro et al., 2015) and (dx.doi.org/10.17504/protocols.io.bv9mn946) with minor modifications. Briefly, cells were dissociated into single cells, filtered through a Flowmi cell strainer and the nuclei were isolated by incubation with lysis buffer (10 mM Tris-HCl pH7.4, 10 mM NaCl, 3 mM MgCl2, 0.1% Tween-20, 0.1% NP-40, 0.01% digitonin and 1% BSA) for 5min at 4C. 20,000 and 40,000 nuclei were isolated, and the DNA was tagmented with Tn5 (Illumina) and amplified using NEBNext® High-Fidelity 2X PCR Master Mix (NEB). The number of cycles was estimated by qPCR. DNA tagmentation efficacy was evaluated with bioanalyzer 2100 (Agilent technologies) and the DNA amounts calculated with Qubit. i5 and i7 primer sequences were obtained from (Mezger et al., 2018). The resulting DNA libraries were sequenced using the NextSeq550 system (Illumina) and about 25 million reads were obtained per sample in duplicate.

### RNA-seq Analysis

Resulting fastq files were mapped to their transcripts and quantified using salmon (Patro et al., 2017) against the genome version mm10. For downstream analyses, Rstudio was used with the packages Deseq2, PCAExplorer and pheatmap (Love et al., 2014; Marini and Binder, 2019). Statistical significance was calculated using a Wald test and corrected using the Benjamini-Hochberg method.

### ATAC-seq Analysis

The raw fast data were analyzed using Basepair software (https://www.basepairtech.com/) with a pipeline that included the following steps. The raw reads were trimmed using fastp to remove low-quality bases from reads (quality < 20) and adapter sequences. The trimmed reads were aligned using Bowtie2 (Langmead and Salzberg, 2012) to UCSC genome assembly mm10. Duplicate reads were removed using Sambamba. Peaks were identified with MACS2 (Gaspar, 2018 preprint) and those overlapping with satellite repeat regions were discarded. For further analyses, a union peak atlas was created from the MACS2 files. Peak intensity for each sample was counted using featureCounts (Liao et al., 2014). HOMER (v4.11) (Heinz et al., 2010) was used for motif analyses. Principal component analyses were done using the R packages DESeq2 (Love et al., 2014) and PCAexplorer (Marini and Binder, 2019).

For motif enrichment analysi, ATAC-seq peaks in the atlas were associated with TF motifs in the CIS-BP database (Weirauch et al., 2014) using FIMO (Grant et al., 2011) of MEME suite (Bailey et al., 2009), under the P value cutoff of 1e-4. We limited the motif analyses to motifs for 236 TFs that are expressed in EpiSCs or DE (RPKM > 5). Normalized counts were obtained using variance-stabilizing transformation in DEseq2. The shift in the cumulative distribution of normalized counts was compared between the subset of the atlas containing the TF motif and the total atlas using one-sided Kolmogorov-Smirnov test.

### Generation of Forebrain Organoids

For forebrain organoid generation, B6129SF1/J EpiSC colonies were detached using Collagenase IV (0.5 u/μL), washed once with PBS^-/-^, and then dissociated into a single cell suspension using Accutase. EpiSCs were then seeded at 1,000 cells/microwell in an Aggrewell plate in EB formation media containing 50 nM chroman-1, 5 μM emricasan, 100 nM LDN, 10 μM SB431542, 100 nM PD173074 and 4 nM LGK974 in N2B27 (B27 without vitamin A) media. After 24h, the EBs were recovered and transferred to the same media but without chroman-1 and emricasan and embedded in Matrigel as previously described (Qian et al., 2018). Briefly, 66.7 μL of EBs + media were mixed with 100uL of Matrigel using wide-bore tips. The mix was added to a non-adherent 6-well plate without touching the edges of the well and incubated at 37C for 30min. After 30 min, 3mL of warm media were added on top. At day 2, the media was changed to plain N2B27 (B27 without vitamin A) with 100 ng/mL Fgf8b and kept for 48 h more. Day 4 EBs were recovered from Matrigel using cell recovery solution (Corning) and incubating for 30 min at 4C. After day the EBs were transferred to a Petri dish and kept in N2B27 (B27 Supplement with vitamin A) with BDNF and GDNF while shaking at 65 rpm (Infors HT, Celltron benchtop shaker). For sparse labeling of neuronal progenitor cells with CytoTune EmGFP Sendai fluorescence reporter, 7.5 μL were added to the 2 mL media in the Aggrewell plate at D0 and kept until D1. The organoids were then imaged for ∼24 hours from d3 to d4.

### Immunofluorescence Staining of Organoids

For organoid staining, the organoids were collected in tubes pre-coated with 1% BSA. The samples were washed twice with PBS^-/-^ and fixed with 4% PFA for 45 min at 4C. Afterwards, they were washed twice with PBS-/- and permeabilized with 0.5% Triton-X from 15 min at 4C. Blocking was performed for 15 min in organoid washing solution (OWS, adapted from (Dekkers et al., 2019), consisting of 0.2% Triton-X, 0.02% SDS and 0.2% BSA in PBS-/-) at 4C. Next, the samples were incubated in OWS with primary antibodies overnight at 4C. On the following day, they were washed 3x for 2h each with OWB while shaking, then incubated overnight at 4C with secondary antibodies and DAPI (1 µM) in OWB. The next day, the organoids were washed 3x with OWB for 6h total while shaking. Finally, the organoids were mounted in DeepClear solution (Celexplorer). The samples were imaged using the confocal Nikon A1RHD25 or Leica SP8 and analyzed with the Imaris software. For live imaging, the organoids were embedded in phenol-free Matrigel and imaged using the TrueLive3D Imager system (Luxendo).

### Single-cell RNA sequencing

Multiple organoids were collected and mixed at day 12, washed with PBS-/- and dissociated with Accutase until a single cell suspension was achieved. Then, the sample was washed once again with PBS-/-. The resulting cell suspension was used for 10X scRNA-seq (10x Genomics, Single Cell 3’ Kit v3.1, dual index) following the manufacturer’s directions. 326M reads were obtained. The resulting fastq files were processed with CellRanger (10X genomics cloud). CellBender was used to eliminate background reads and other artifacts (Fleming et al., 2019 preprint), and, any cell with more than 10% mitochondrial reads was excluded. The Seurat package (v4) was used for downstream analyses (Hao et al., 2021). Cluster annotation was performed using the cited literature and the Allen Developing Mouse Brain Atlas, stages E15.5 and E18.5. VoxHunt analyses were performed as described in (Fleck et al., 2021).

## Supporting information

Supplementary experimental methods

## ACKNOWLEDGEMENTS

This study would not have been possible without the core facilities at MSKCC. We would like to specifically thank Dr. Yas Furuta (Mouse Genetics Core facility, MSKCC), Murray Tipping and Vitaly Boyko (Molecular Cytology Core, MSKCC), Dr. Richard Koche (Epigenetics Innovation Lab, MSKCC), and members of the Integrated Genomics Operation Core (MSKCC) for their advice and assistance with experiments and data analysis throughout the course of this project. In addition, we would like to thank Dr. James Muller (Director of Developmental Biology Program’s Microscope Cluster) for assistance with light-sheet microscopy and advance about imaging organoids and Dr. Ronan Chaligne (Single Cell Research Initiative, MSKCC) for advice and assistance with single cell RNA-sequencing analysis.

We would like to thank Dr. Laura Reinholdt and Anne Czechanski (Jackson Labs) for sharing PWK/PhJ embryonic stem cell lines, Dr. Christopher Baker and Dr. Candice Byers (Jackson Labs) for sharing C57Bl/6J embryonic stem cell lines, and Dr. Matthias Stadtfeld (Weill Cornell Medical College) for sharing B6129SF1/J embryonic stem cell lines.

We would like to thank our colleagues Dr. Anna Katerina Hadjantonakis, Dr. Danwei Huangfu, Dr. Lorenz Studer, Dr. Nan Yang, Dr. Matthias Stadtfeld, Dr. Eftychia Apostolou, Dr. Sophie Morgani, Dr. Clayton Schwarz, Dr. Gabriele Ciceri, Dr. Ryan Walsh, Bess Rosen (Huangfu Lab), Renhe Luo (Huangfu lab), Dapeng Yang (Huangfu Lab) for helpful discussions and advice throughout the course of the study. Figure panels 2C, 3F and 4A were created using Biorender.com

We would like to thank Dr. Nan Yang (Mount Sinai), Dr. Matthias Stadtfeld, Dr. Timothy Cherry for helpful comments on the manuscript.

This study was made possible by financial support from: Startup funding from the Sloan Kettering Institute and Memorial Sloan Kettering Cancer Center (T.V.) and the Josie Robertson Investigator Program (T.V.), a NARSAD Young Investigator Grant from the Brain & Behavior Research Foundation (#28756) (T.V.)., Tri-Institutional Stem Cell Initiative (Starr Foundation), and a training award from NYSTEM (contract # C32559GG) and the Center for Stem Cell Biology at MSK (D.M.C.).

## AUTHOR CONTRIBUTIONS

Conceptualization: T.V., D.M.C., E.K.C.

Methodology: D.M.C., E.K.C., R.A.G., T.V.

Investigation: D.M.C., E.K.C, R.A.G., M.I., Y.L., H.C., J.K., T.V.

Data curation: D.M.C.

Writing: D.M.C., E.K.C., R.A.G., T.V.

Funding Acquisition: T.V.

Supervision: T.V.

## DATA AVAILABILITY

Source data and processed files are available in the Gene expression Omnibus under the accession numbers GSE189869 (RNA and ATAC-seq datasets) and GSE189870 (scRNA-seq).

## DECLARATION OF INTERESTS

The authors have no competing interests to declare.

### SUPPLEMENTARY FIGURE LEGENDS

**Supplementary Figure 1. Systematic optimization of conditions for EpiSC-DE differentiation.** (**A**) Overview of state-of-the-art differentiation protocols for human (purple) and mouse (green) PSCs. Further details of each study are provided in **Supplementary Table 1**). (**B-C**) Results of DE protocol optimization experiments. For each condition, immunofluorescence staining was performed following 40 h of differentiation (end of Stage 2). Immunofluorescence images for the indicated antibodies were quantified using CellProfiler. Markers of DE (FOXA2) and mesoderm (CDX2) are shown. (**B**) Summary of results from anterior primitive streak (Stage 1) optimization experiments (n=2 technical replicates). After the indicated conditions for Stage 1, all conditions were exposed to the same Stage 2 conditions, and then quantified. (**C**) Summary of results for optimization of DE commitment (Stage 2) conditions (n=2 technical replicates). For Stage 1, the optimal conditions identified in (**B**) were used, and then cells were switched into the indicated conditions for Stage 2.

**Supplementary Figure 2. Extended characterization of EpiSC-derived definitive endoderm.** (**A**) Euclidean distance between samples based on ATAC-seq (n=4 per condition, left) and RNA-seq (n=5 for EpiSCs and n=4 for MACs-sorted DE samples, right). (**B**) PCA plot of bulk chromatin accessibility (left) and gene expression (right) of the same C57Bl/6J EpiSC and DE samples as (**A**). (**C**) Unbiased clustered heatmap of the top variable genes as detected by RNA-seq. (**D**) Expression of genes previously identified as DE markers in single cell gene expression studies of hPSC-derived DE and DE isolated from E7.5 embryos (Genga et al., 2019; Nowotschin et al., 2019).

**Supplementary Figure 3. Immunostainings of EpiSC-derived definitive endoderm.** (**A**) Representative immunofluorescence images from DE differentiations (n = 3 biological replicates) for a panel of DE markers (FOXA2, GATA6, SOX17, OTX2, EOMES), pluripotency markers (OCT4), and primitive streak/mesoderm markers (T, CDX2). (**B**) Representative immunofluorescence images from DE differentiations (n = 3 biological replicates) performed using EpiSCs (+WI) cultured in feeder-free conditions prior to starting the differentiation protocol. Scale bar = 50 μm.

**Supplementary Figure 4: Optimization of conditions for forebrain organoid generation from mouse EpiSCs.** (**A**) Brightfield images of d1 organoids in Aggrewell wells. EBs were formed in the presence of Thiazovivin or chroman 1 + emricasan (CE), or the recently published CEPT cocktail (chroman 1, emricasan, polyamine mix, trans-ISRIB) (Chen et al., 2021). Scale bar = 100 μm. (**B**) Confocal immunofluorescence image of a day 4 EB/organoid. Scale bar = 25 μm.

**Supplementary Figure 5: EpiSC-derived brain organoids generate prethalamic and cortical-like neuronal populations.** (**A,B**) Confocal immunofluorescence images of d8 organoids stained with antibodies against classical cortical markers (TBR1, TBR2, RELN and PAX6), which are also found in other regions of the brain such as the hippocampus, the cortical hem and the prethalamus, and thalamic markers (TCF7L2). Scale bar = 100 μm. (**C-D**) Confocal immunofluorescence of d8 organoids from C57Bl/6J background. Scale bar = 100 μm. (**E**) Confocal immunofluorescence image of a d12 organoid stained using antibodies against the classical upper-layer cortical marker SATB2. Scale bar = 100 μm. (**F-G**) Confocal immunofluorescence images of d12 organoids. Scale bar = 100 μm.

**Supplementary Figure 6: scRNA-seq analyses on day 12 organoids reveals three main cell types**. (**A**) Heatmap of the top 10 genes that define each cluster. Abbreviations: VZ = ventricular zone, MZ = mantle zone. (**B**) Expression profile, across clusters, of several prethalamic (*Sp9, Arx, Pax6, Meis2, Ptprd, Islr2*) and axonal (*L1cam, Nrcam*) markers. (**C**) Confocal immunofluorescence image of a day 12 organoid. Scale bar = 30 µm.

**Supplementary Table 1: Summary of previous mESC, mEpiLC, and hPSC directed differentiation protocols (related to Figure 2).** References used to generate the plot in Fig. 2A (Borowiak et al., 2009; Chen et al., 2013; D’Amour et al., 2005; Diekmann et al., 2019; Green et al., 2011; Kinoshita et al., 2020; Korostylev et al., 2017; Li et al., 2011; Loh et al., 2014; Mfopou et al., 2014; Morrison et al., 2016; Morrison et al., 2008; Mou et al., 2012; Mulas et al., 2017; Ortmann et al., 2020; Sherwood et al., 2011; Shi et al., 2017).

**Supplementary Table 2: List of antibodies used. Supplementary Table 3: List of reagents used. Supplementary Table 4: Mouse embryonic stem cell lines**

**Supplementary Table 5: EpiSC lines used in brain organoid experiments. Supplementary Table 6: Complete UMAPs, related to figure 6**.

**Supplementary Video 1. Day 4 organoid stained with antibodies against ZO-1 (green), PH3 (purple) and Nestin (red).**

**Supplementary Video 2. Live imaging of radial glial-like cells in a d3 organoid.** emGFP Sendai virus was added to the EB formation media during the EB formation period to sparsely label developing neuroepithelial progenitor cells. The organoid was then embedded in Matrigel and imaged with a light sheet microscope from day 3 to day 4. The apical membrane is at the bottom of the video and the basal lamina on top.

**Supplementary Experimental Methods 1: Step-by-step protocols for directed differentiation experiments**

