## Supplementary figures and tables for "Rapid and robust directed differentiation of mouse epiblast stem cells into definitive endoderm and forebrain organoids"

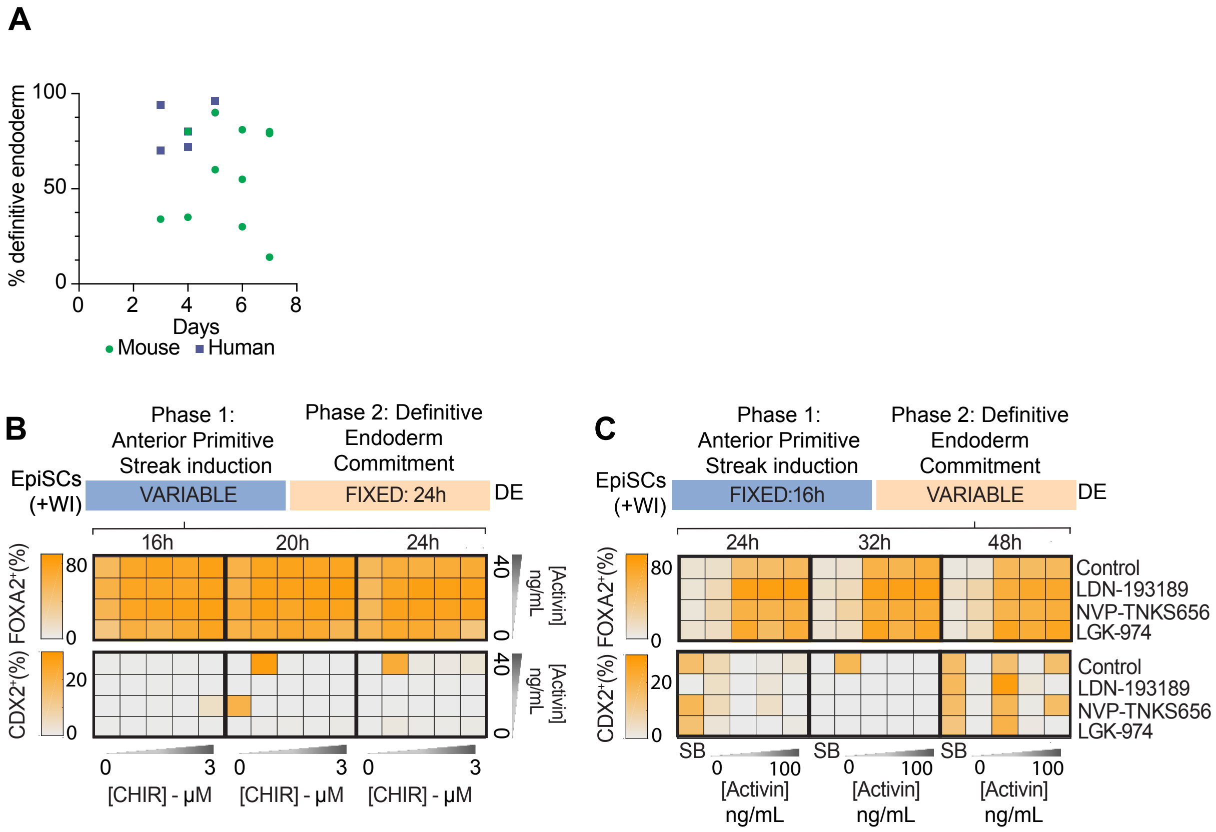
A
% definitive endoderm
100
50
0
0 2 4 6 8
Days
Mouse
Human
B
Phase 1: Anterior Primitive Streak induction
Phase 2: Definitive Endoderm Commitment
EpiSCs (+WI)
VARIABLE
FIXED: 24h
DE
16h
20h
24h
FOXA2⁺(%)
80
0
CDX2⁺(%)
20
0
40
0
[Activin] ng/mL
0 3
[CHIR] - µM
C
Phase 1: Anterior Primitive Streak induction
Phase 2: Definitive Endoderm Commitment
EpiSCs (+WI)
FIXED:16h
VARIABLE
DE
24h
32h
48h
Control
LDN-193189
NVP-TNKS656
LGK-974
SB
0 100
[Activin] ng/mL

**Supplementary Figure 1. Systematic optimization of conditions for EpiSC-DE differentiation.** (**A**) Overview of state-of-the-art differentiation protocols for human (purple) and mouse (green) PSCs. Further details of each study are provided in **Supplementary Table 1**). (**B-C**) Results of DE protocol optimization experiments. For each condition, immunofluorescence staining was performed following 40 h of differentiation (end of Stage 2). Immunofluorescence images for the indicated antibodies were quantified using CellProfiler. Markers of DE (FOXA2) and mesoderm (CDX2) are shown. (**B**) Summary of results from anterior primitive streak (Stage 1) optimization experiments (n=2 technical replicates). After the indicated conditions for Stage 1, all conditions were exposed to the same Stage 2 conditions, and then quantified. (**C**) Summary of results for optimization of DE commitment (Stage 2) conditions (n=2 technical replicates). For Stage 1, the optimal conditions identified in (**B**) were used, and then cells were switched into the indicated conditions for Stage 2.

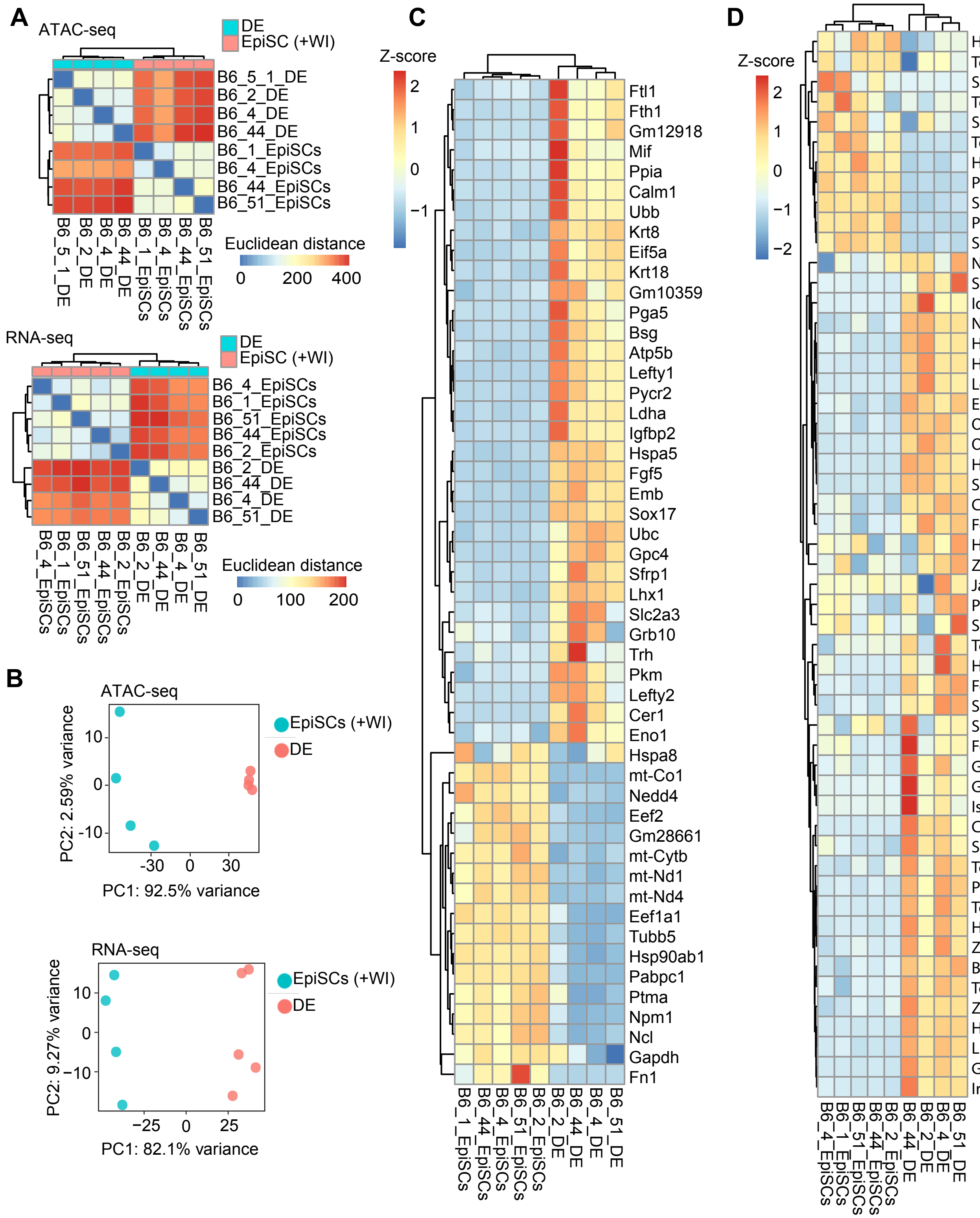

A
ATAC-seq
DE
EpiSC (+WI)
B6_5_1_DE
B6_2_DE
B6_4_DE
B6_44_DE
B6_1_EpiSCs
B6_4_EpiSCs
B6_44_EpiSCs
B6_51_EpiSCs
Euclidean distance
0 200 400
RNA-seq
DE
EpiSC (+WI)
B6_4_EpiSCs
B6_1_EpiSCs
B6_51_EpiSCs
B6_44_EpiSCs
B6_2_EpiSCs
B6_2_DE
B6_44_DE
B6_4_DE
B6_51_DE
Euclidean distance
0 100 200
B
ATAC-seq
EpiSCs (+WI)
DE
PC2: 2.59% variance
PC1: 92.5% variance
RNA-seq
EpiSCs (+WI)
DE
PC2: 9.27% variance
PC1: 82.1% variance
C
Z-score
Ftl1
Fth1
Gm12918
Mif
Ppia
Calm1
Ubb
Krt8
Eif5a
Krt18
Gm10359
Pga5
Bsg
Atp5b
Lefty1
Pycr2
Ldha
Igfbp2
Hspa5
Fgf5
Emb
Sox17
Ubc
Gpc4
Sfrp1
Lhx1
Slc2a3
Grb10
Trh
Pkm
Lefty2
Cer1
Eno1
Hspa8
mt-Co1
Nedd4
Eef2
Gm28661
mt-Cytb
mt-Nd1
mt-Nd4
Eef1a1
Tubb5
Hsp90ab1
Pabpc1
Ptma
Npm1
Ncl
Gapdh
Fn1
B6_1_EpiSCs
B6_44_EpiSCs
B6_4_EpiSCs
B6_51_EpiSCs
B6_2_EpiSCs
B6_2_DE
B6_44_DE
B6_4_DE
B6_51_DE
D
Z-score
Hmga1
Tead2
Shh
Tgif1
Smarce1
Tdgf1
Hmga2
Pbx1
Sox11
Pou5f1
Smarcc1
Nfyb
Sp5
Id3
Nodal
Hmgb3
Hes1
Lefty1
E2f5
Otx2
Ovol2
Hhex
Sox17
Cxcr4
Foxa2
Hmgb1
Zfp664
Jarid2
Peg3
Sall1
Tcf4
Hesx1
Foxa1
Sall4
Six1
Foxd4
Gata4
Gata3
Isl1
Cer1
Sox4
Tcf7l1
Prdm1
Tcf7l2
Hdac6
Zic3
Brd3
Tcf3
Zfp422
Hnf1b
Lhx1
Gata6
Irx3
B6_4_EpiSCs
B6_1_EpiSCs
B6_51_EpiSCs
B6_44_EpiSCs
B6_2_EpiSCs
B6_44_DE
B6_2_DE
B6_4_DE
B6_51_DE

**Supplementary Figure 2. Extended characterization of EpiSC-derived definitive endoderm.** (**A**) Euclidean distance between samples based on ATAC-seq (n=4 per condition, left) and RNA-seq (n=5 for EpiSCs and n=4 for MACs-sorted DE samples, right). (**B**) PCA plot of bulk chromatin accessibility (left) and gene expression (right) of the same C57Bl/6J EpiSC and DE samples as (**A**). (**C**) Unbiased clustered heatmap of the top variable genes as detected by RNA-seq. (**D**) Expression of genes previously identified as DE markers in single cell gene expression studies of hPSC-derived DE and DE isolated from E7.0 embryos (Genga et al., 2019; Nowotschin et al., 2019).

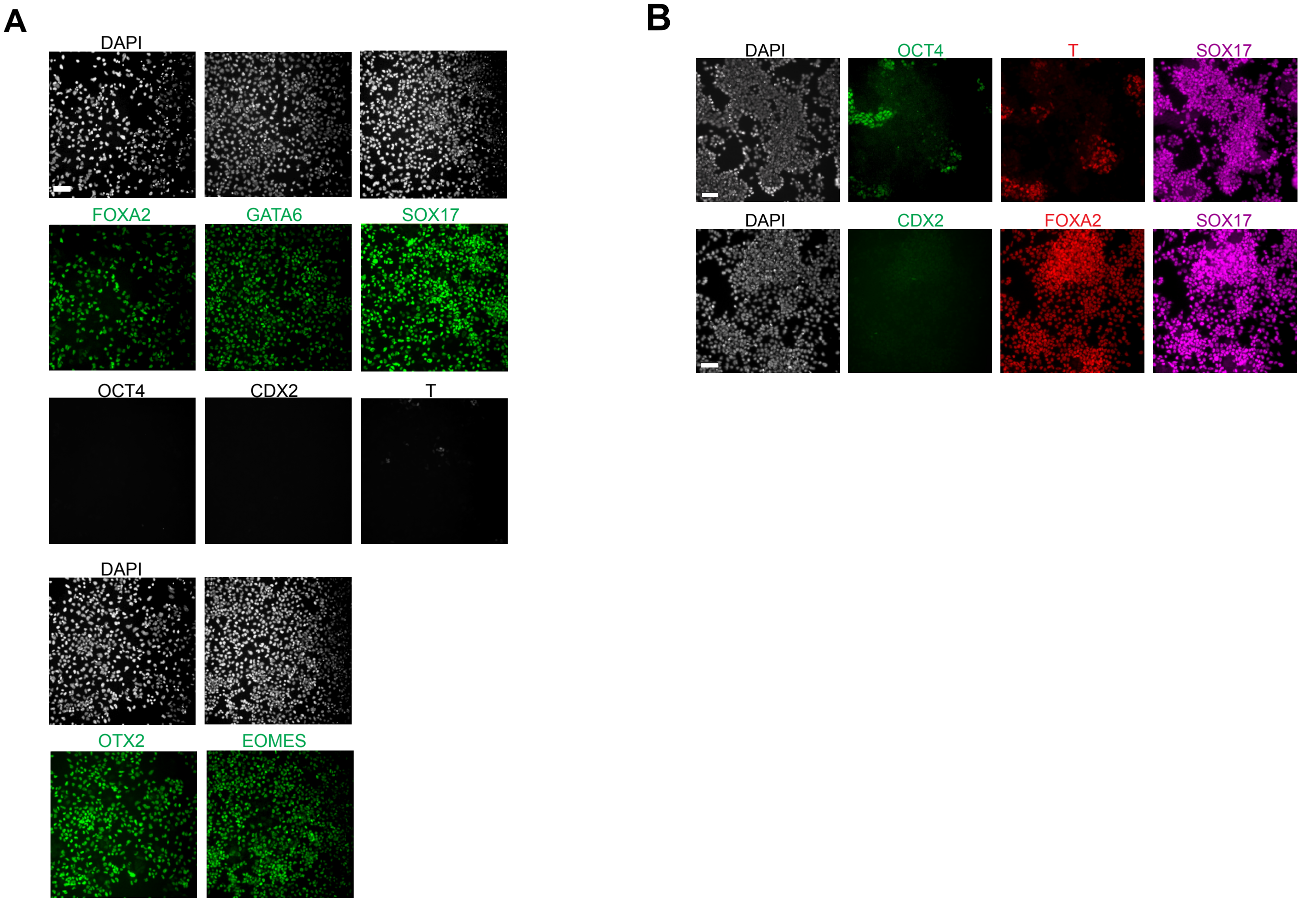
A
DAPI
FOXA2
GATA6
SOX17
OCT4
CDX2
T
DAPI
OTX2
EOMES
B
DAPI
OCT4
T
SOX17
DAPI
CDX2
FOXA2
SOX17

**Supplementary Figure 3.** **Immunostainings of EpiSC-derived definitive endoderm.** (**A**) Representative immunofluorescence images from DE differentiations for a panel of DE markers (FOXA2, GATA6, SOX17, OTX2, EOMES), pluripotency markers (OCT4), and primitive streak/mesoderm markers (T, CDX2). (**B**) Immunofluorescence images from feeder-free derived DE for SOX17, FOXA2, T, OCT4 and CDX2). All images represent the average results after 4 or more experiments, with at least 3 biological replicates each. Scale bar = 50 μm.

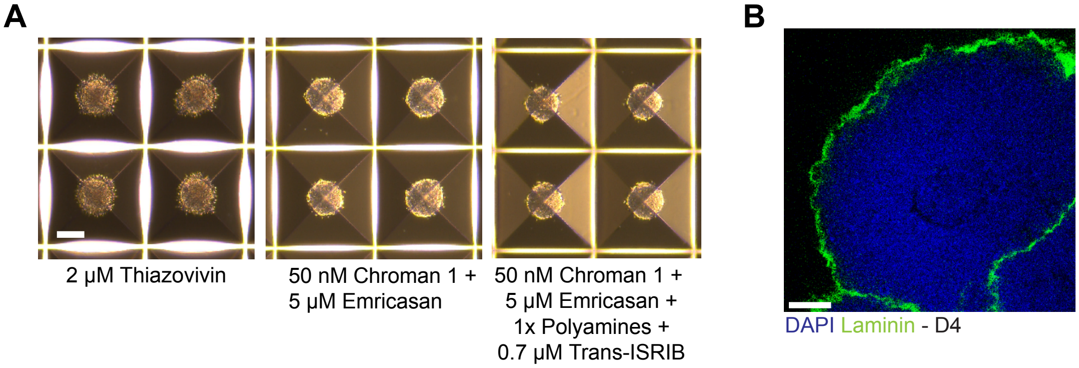

A
2 μM Thiazovivin
50 nM Chroman 1 +
5 μM Emricasan
50 nM Chroman 1 +
5 μM Emricasan +
1x Polyamines +
0.7 μM Trans-ISRIB
B
DAPI Laminin - D4

**Supplementary Figure 4: Optimization of conditions for forebrain organoid generation from mouse EpiSCs.** (**A**) Brightfield images of d1 organoids in Aggrewell wells. EBs were formed in the presence of Thiazovivin or chroman 1 + emricasan (CE), or the recently published CEPT cocktail (chroman 1, emricasan, polyamine mix, trans-ISRIB) (Chen et al., 2021). Scale bar = 100 μm. (**B**) Confocal immunofluorescence image of a day 4 EB/organoid. Scale bar = 25 μm.

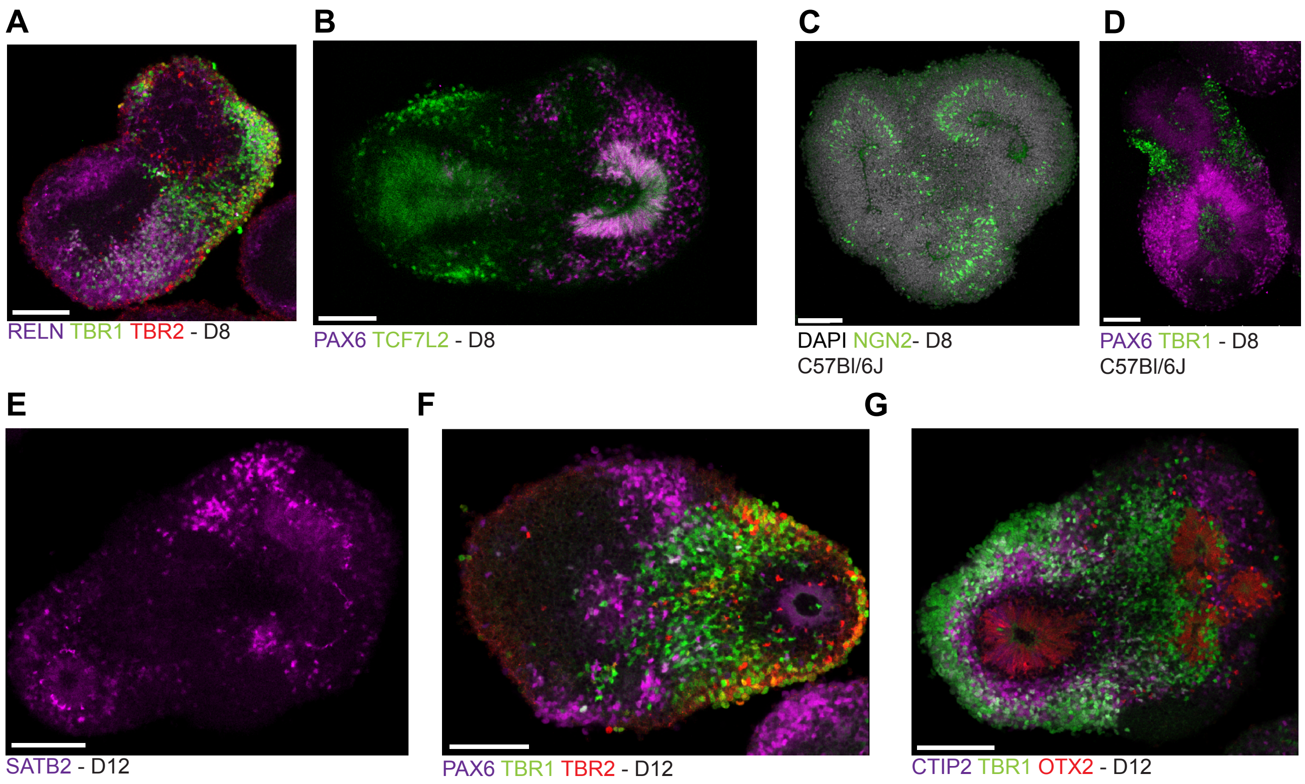
A
RELN TBR1 TBR2 - D8
B
PAX6 TCF7L2 - D8
C
DAPI NGN2- D8
C57Bl/6J
D
PAX6 TBR1 - D8
C57Bl/6J
E
SATB2 - D12
F
PAX6 TBR1 TBR2 - D12
G
CTIP2 TBR1 OTX2 - D12

**Supplementary Figure 5: EpiSC-derived brain organoids generate prethalamic and cortical-like neuronal populations.** (**A,B**) Confocal immunofluorescence images of d8 organoids stained with antibodies against classical cortical markers (TBR1, TBR2, RELN and PAX6), which are also found in other regions of the brain such as the hippocampus, the cortical hem and the prethalamus, and thalamic markers (TCF7L2). Scale bar = 100 µm. (**C-D**) Confocal immunofluorescence of d8 organoids from C57Bl/6J background. Scale bar = 100 µm. (**E**) Confocal immunofluorescence image of a d12 organoid stained using antibodies against the classical upper-layer cortical marker SATB2. Scale bar = 100 µm. (**F-G**) Confocal immunofluorescence images of d12 organoids. Scale bar = 100 µm.

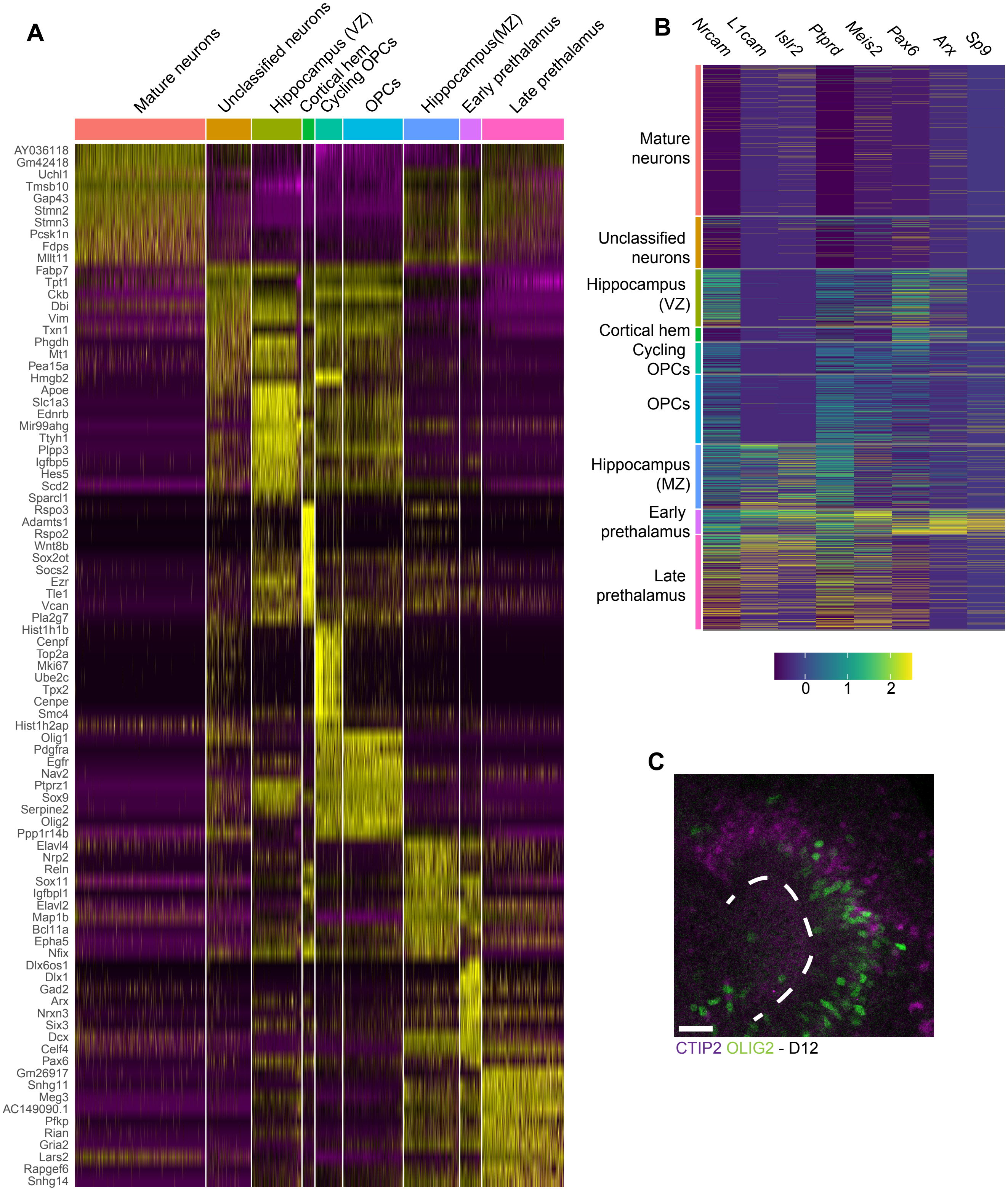

A
Mature neurons
Unclassified neurons
Hippocampus (VZ)
Cortical hem
Cycling OPCs
OPCs
Hippocampus(MZ)
Early prethalamus
Late prethalamus
AY036118
Gm42418
Uchl1
Tmsb10
Gap43
Stmn2
Stmn3
Pcsk1n
Fdps
Mllt11
Fabp7
Tpt1
Ckb
Dbi
Vim
Txn1
Phgdh
Mt1
Pea15a
Hmgb2
Apoe
Slc1a3
Ednrb
Mir99ahg
Ttyh1
Plpp3
Igfbp5
Hes5
Scd2
Sparcl1
Rspo3
Adamts1
Rspo2
Wnt8b
Sox2ot
Socs2
Ezr
Tle1
Vcan
Pla2g7
Hist1h1b
Cenpf
Top2a
Mki67
Ube2c
Tpx2
Cenpe
Smc4
Hist1h2ap
Olig1
Pdgfra
Egfr
Nav2
Ptprz1
Sox9
Serpine2
Olig2
Ppp1r14b
Elavl4
Nrp2
Reln
Sox11
Igfbpl1
Elavl2
Map1b
Bcl11a
Epha5
Nfix
Dlx6os1
Dlx1
Gad2
Arx
Nrxn3
Six3
Dcx
Celf4
Pax6
Gm26917
Snhg11
Meg3
AC149090.1
Pfkp
Rian
Gria2
Lars2
Rapgef6
Snhg14
B
Nrcam
L1cam
Islr2
Ptprd
Meis2
Pax6
Arx
Sp9
Mature neurons
Unclassified neurons
Hippocampus (VZ)
Cortical hem
Cycling OPCs
OPCs
Hippocampus (MZ)
Early prethalamus
Late prethalamus
0
1
2
C
CTIP2 OLIG2 - D12

**Supplementary Figure 6: scRNA-seq analyses on day 12 organoids reveals three main cell types**. (**A**) Heatmap of the top 10 genes that define each cluster. Abbreviations: VZ = ventricular zone, MZ = mantle zone. (**B**) Expression profile, across clusters, of several prethalamic (*Sp9, Arx, Pax6, Meis2, Ptprd, Islr2*) and axonal (*L1cam, Nrcam*) markers. (**C**) Confocal immunofluorescence image of a day 12 organoid. Scale bar = 30 μm.

| PMID | Organism | Days | Efficiency | Journal | Published year |
|---|---|---|---|---|---|
| 26675138 | Mouse | 7 | 80 | The Embo journal | 2016 |
| 33271069 | Mouse | 3 | FS cells: 34%, EpiSCs 4% | Cell Stem Cell | 2021 |
| 28669603 | Mouse | 5-6 | 60% | Stem Cell Reports | 2017 |
| 24239964 | Mouse | 6 | 54.92 | Stem cell research | 2014 |
| 19341624 | Mouse | 6 | 81 | Cell stem cell | 2009 |
| 23293299 | Mouse | 7 | 79 | Development | 2013 |
| 28702321 | Mouse | 4 | 35 | Molecular metabolism | 2017 |
| 32795399 | Mouse | 6 | 30 | Cell stem cell | 2020 |
| 18940732 | Mouse | 7 | 14 | Cell stem cell | 2008 |
| 21400570 | Mouse | 4 | 80 | Cellular Biochemistry | 2011 |
| 22482504 | Mouse | 5 | >90% | Cell Stem Cell | 2012 |
| 21854845 | Mouse | 5 | >90% | Mechanisms of Development | 2011 |
| 24412311 | Human | 3 | 94 | Cell stem cell | 2014 |
| 28196600 | Human | 4 | 72 | Cell stem cell | 2017 |
| 16258519 | Human | 4 | 80 | Nature biotechnology | 2005 |
| 30700818 | Human | 3 | 70 | nature scientific reports | 2019 |
| 21358635 | Human | 5 | 96 | Nature biotechnology | 2011 |

**Supplementary Table 1: Summary of previous mESC, mEpiLC, and hPSC directed differentiation protocols (related to Figure 2).** References used to generate the plot in Fig. 2A (Borowiak et al., 2009; Chen et al., 2013; D'Amour et al., 2005; Diekmann et al., 2019; Green et al., 2011; Kinoshita et al., 2020; Korostylev et al., 2017; Li et al., 2011; Loh et al., 2014; Mfopou et al., 2014; Morrison et al., 2016; Morrison et al., 2008; Mou et al., 2012; Mulas et al., 2017; Ortmann et al., 2020; Sherwood et al., 2011; Shi et al., 2017).

| Antibody | Concentration | Manufacturer | Cat.No. |
|---|---|---|---|
| Alexa Fluor 488 Anti-Human SOX17 conjugated antibody | 5uL/10^6 cells | R&D Systems, Inc. | 614013 |
| Alexa Fluor 647 Anti-mouse CD184 conjugated antibody | 0.5ug/10^6 cells | BioLegend | 146504 |
| Anti-CDX2 | 1/250 | Abcam | Ab157524 |
| Anti-CTIP2 | 1/200 | Abcam | Ab18465 |
| Anti-FOXA2 | 1/500 | Abcam | Ab108422 |
| Anti-GATA4 | 1/200 | Santa Cruz | sc-25310 |
| Anti-GATA6 | 1/500 | Cell signaling technology | 5851S |
| Anti-GFAP | 1/1000 | Invitrogen | 13-0300 |
| Anti-Human SOX17 | 1/200 | R&D Systems | AF1924 |
| Anti-Human/Mouse BRACHYURY antibody | 1/200 | R&D systems | AF2085 |
| Anti-LAMININ | 1/1000 | Sigma | L9393 |
| Anti-NESTIN | 1/500 | Aves | NES |
| Anti-NEUROGENIN-2 | 1/200 | R&D Systems | MAB3314-SP |
| Anti-OCT4 | 1/500 | Invitrogen | 701756 |
| Anti-OLIG2 | 1/500 | Abcam | Ab109186 |
| Anti-OTX1+OTX2 | 1/500 | Abcam | Ab21990 |
| Anti-OTX2 | 1/500 | R&D Systems | AF1979 |
| Anti-PAX6 | 1/200 | BD Biosciences | 561462 |
| Anti-PDX1 | 1/100 | Abcam | Ab47308 |
| Anti-PH3 | 1/500 | Cell Signaling | 9706S |
| Anti-REELIN | 1/200 | Millipore | MAB5366 |
| Anti-SATB2 | 1/200 | Abcam | Ab51502 |
| Anti-SOX1 | 1/200 | R&D Systems | AF3369 |
| Anti-SOX2 | 1/500 | Sigma | AB5603 |
| Anti-TBR1 | 1/500 | Abcam | Ab31940 |
| Anti-TBR2 (for brain organoids) | 1/500 | Millipore | AB15894 |

| | | | |
|---|---|---|---|
| Anti-TBR2/ EOMES (for 2D staining) | 1/500 | Abcam | Ab23345 |
| Anti-TCF7L2 | 1/200 | Cell Signaling | 2569 |
| Anti-TUJ1 | 1/1000 | BioLegend | 801213 |
| Anti-ZO-1 | 1/500 | Thermo | 617300 |
| Live/dead zombie UV staining | 1uL/10^6 cells | Invitrogen | L34962 |
| DAPI | 300nM (ICC)<br>1uM (organoid) | Invitrogen | D1306 |
| Donkey anti Rabbit IgG Secondary Antibody, Alexa Fluor 488 | 1/500 | Invitrogen | A-21206 |
| Donkey anti-Goat highly cross-adsorbed secondary antibody, Alexa Fluor Plus 488 | 1/500 | Invitrogen | A-32814 |
| Donkey anti Rat IgG Highly Cross Adsorbed Secondary Antibody, Alex Fluor 488 | 1/500 | Invitrogen | A-21208 |
| Donkey anti Mouse IgG Highly Cross Adsorbed Secondary Antibody, Alexa Fluor 488 | 1/500 | Invitrogen | A-21202 |
| Goat anti-Chicken IgY Secondary Antibody, Alexa Fluor 555 | 1/500 | Invitrogen | A-21437 |
| Donkey anti Mouse IgG Secondary Antibody, Alexa Fluor 555 | 1/500 | Invitrogen | A-31570 |
| Donkey anti Rabbit IgG Secondary Antibody, Alexa Fluor 555 | 1/500 | Invitrogen | A-31572 |
| Donkey anti Goat IgG Cross Adsorbed Secondary Antibody Alexa Fluor 555 | 1/500 | Invitrogen | A-21432 |
| Chicken anti-Rat IgG Cross Adsorbed Secondary Antibody, Alexa Fluor 647 | 1/500 | Invitrogen | A-21472 |

| | | | |
|---|---|---|---|
| Chicken ant Goat IgG Secondary Antibody, Alexa Fluor 647 | 1/500 | Invitrogen | A-21469 |
| Donkey anti Mouse IgG Secondary Antibody, Alexa Fluor 647 | 1/500 | Invitrogen | A-31571 |
| Donkey anti Rabbit IgG Highly Cross Adsorbed Secondary Antibody, Alexa Fluor 647 | 1/500 | Invitrogen | A-31573 |

**Supplementary Table 2: List of antibodies used.**

| **Reagent** | **Manufacturer** | **Reference #** |
|---|---|---|
| 2-mercaptoethanol | Gibco | 21985023 |
| Accutase | Gibco | A1110501 |
| Activin A | Peprotech | 120-14P |
| Aggrewell plate | STEMCELL | 34415 |
| Anti-Rat IgG Microbeads | Miltenyi Biotec | 130-048-502 |
| Apo-transferrin | InVitra | 777TRF029 |
| B27 supplement | ThermoScientific | 17504044 |
| B27 supplement (without vitamin A) | Gibco | 12587010 |
| BDNF | PeproTech | 450-02 |
| BenchMark Fetal bovine serum | Gemini | 100-106 |
| bFGF | Gibco | 12587010 |
| BMP-7 | PeproTech | 120-03P |
| Bovine Serum Albumin | Sigma | A2153 |
| Cell recovery solution | Corning | 354253 |
| Chemically Defined Lipid Concentrate | Gibco | 11905031 |
| CHIR99201 | Tocris Bioscience | 252917-06-9 |
| Chroman 1 | MedChem Express | HY-15392 |
| Chromium Next GEM Chip G Single Cell Kit, 48 rxns | 10X genomics | 1000120 |
| Chromium Next GEM Single Cell 3' Kit v3.1, 16 rxns | 10X genomics | 1000268 |
| CytoTune emGFP Sendai fluorescence reporter | ThermoScientific | A16519 |
| Collagenase Type IV | Gibco | 17104019 |
| DAPI | Thermo Scientific | D1306 |
| DeepClear | Celexplorer | DC-201 |
| Digitonin | Sigma | D-141 |
| DMEM with glutamax | Gibco | 10564029 |
| DMEM/F12 | Gibco | 11320082 |
| Dual Index Kit TT Set A 96 rxns | 10x genomics | 1000215 |
| eBioscience Fixation/Permeabilization Concentrate | Invitrogen | 00-5123 |

| | | |
|---|---|---|
| eBioscience Fixation/Permeabilization Dilent | Invitrogen | 00-5223-56 |
| eBioscience Permeabiliation buffer (10X) | Invitrogen | 00-8333 |
| EDTA (0.5 M), pH 8.0, RNase-free | Invitrogen | AM9261 |
| Emricasan | Selleckchem | S7775 |
| ESGRO LIF | Sigma | ESG1107 |
| F12 with GlutaMAX | Gibco | 31765035 |
| Fetal bovine serum | Gemini | 100-106 |
| Fgf-8b protein, CF | R&D Systems | 423-F8-025/CF |
| Fibronectin | Sigma | FC010 |
| Fixation/permeabilization concentrate | Invitrogen | 00-5223-43 |
| Fixation/permeabilization diluent | Invitrogen | 00-5223 |
| Flowmi cell strainer | Bel-Art | 136800040 |
| GDNF | PeproTech | 450-10 |
| Glutamax | Gibco | 35050061 |
| Heat stable recombinant human bFGF | Gibco | PHG0360 |
| Hyclone fetal bovine serum | Cytiva | SH3007003 |
| IMDM | Gibco | 12440053 |
| Insulin | Sigma | 91077C |
| KAPA Hyper Prep Kit | Kapa Biosystems | KK8504 |
| KingFisher Flex Magnetic Particle Processor | Thermo Scientific | 5400630 |
| Knockout serum replacement | Gibco | 10828028 |
| Laminin from Engelbreth-Holm-Swarm murine sarcoma basement membrane | Sigma | L2020 |
| LDN-193189 | Stemgent | 04-0074 |
| LGK-974 | Selleck Chemicals | S7143 |
| LY294002 | Selleckchem | S1105 |
| MACS multistand | Miltenyi Biotech | 130-042-303 |
| MagMAX *mir*Vana Total RNA Isolation Kit | Thermo Scientific | A27828 |
| Matrigel | Corning | 354230 |
| Monothioglycerol | Sigma | M6145 |
| MS column | Miltenyi Biotech | 130-122-727 |
| N2 supplement | Gibco | 17502048 |

| | | |
|---|---|---|
| NEBNext® High-Fidelity 2X PCR Master Mix | NEB | M0541L |
| Neurobasal Media | Gibco | 21103049 |
| Non-adherent 6 well plate | Corning | 3471 |
| Non-essential amino acids | Gibco | 11140050 |
| NP-40 detergent | Sigma | 74385 |
| NVP-TNKS656 | Selleck Chemicals | S7238 |
| OctoMACS separator | Miltenyi Biotech | 130-042-109 |
| PD0325901 | Tocris | 4192 |
| PD173074 | STEMCELL Technologies | 72164 |
| Penicllin/streptomycin (100X) | Gibco | 15140122 |
| Permeabilization buffer | Invitrogen | 00-8333 |
| PiK-90 | Sigma Aldrich | 528117 |
| Polyvinyl Alcohol | Sigma | 341584 |
| Putrescine | Sigma | P5780 |
| rhLaminin-521 | Gibco | A29249 |
| SB-431542 | Tocris | 1614 |
| SMART-Seq v4 Ultra Low Input RNA Kit | Clonetech | 63488 |
| Sodium pyruvate | Gibco | 11360070 |
| Spermidine | Sigma | S2626 |
| Spermine | Sigma | S4264 |
| 16% Formaldehyde (w/v), Methanol-free | Thermo Scientific | 28908 |
| Thiazovivin | Sigma | SML1045 |
| Tn5 enzyme and tagmentation buffer | Illumina | 20034198 |
| Trans-isrib | Tocris | 5284 |
| Tri Reagent | Sigma | T9424 |
| TruSeq Stranded mRNA LT Kit | Illumina | RS-122-2102 |
| TrypLE | Thermo Scientific | 12605028 |
| Triton X-100 | Sigma-Aldrich | 9002-93-1 |
| Tween-20 | Boston Bioproducts | P-934 |

**Supplementary Table 3: List of reagents used.**

| Name | B6.129_17 | B6.129_9 | B6.129_4 | DBA/2J_55 | C57Bl/6J_44 | C57Bl/6J_51 | C57Bl/6J_1 | C57Bl/6J_4 | C57Bl/6J_2 | PWK/PhJ_AC705 | PWK/PhJ_AC698 | PWK/PhJ_AC695 | PWK/PhJ_AC699 |
|---|---|---|---|---|---|---|---|---|---|---|---|---|---|
| Background | B6129SF1/J | B6129SF1/J | B6129SF1/J | DBA/2J | C57Bl/6J | C57Bl/6J | C57Bl/6J | C57Bl/6J | C57Bl/6J | PWK/PhJ | PWK/PhJ | PWK/PhJ | PWK/PhJ |
| Received from | Matthias Stadtfeld, PhD - Weill Cornell | | | Christopher L. Baker, Ph.D. - The Jackson Laboratory | | | inhouse, through IVF | | | Laura Reinholdt, Ph.D. - The Jackson Laboratory | | | |
| Culture conditions (mESCs) | On gelatin-coated dishes with irradiated mouse embryonic fibroblast feeder cells | | | | | | | | | | | | |
| Media composition (mESCs) | Serum/LIF media: DMEM with glutamax, 1% nonessential amino acids, 1% sodium pyruvate, 1% penicillin-streptomycin, 0.1% 2-mercaptoethanol, 10% fetal bovine serum and 1% ESGRO LIF. | | | Serum/LIF media: DMEM with glutamax, 1% nonessential amino acids, 1% sodium pyruvate, 1% penicillin-streptomycin, 0.1% 2-mercaptoethanol, 10% fetal bovine serum, 1% ESGRO LIF, 3 uM CHIR99201 and 1 uM PD0325901. | | | | | | | | | |
| Culture conditions (mEpiSCs) | On gelatin-coated dishes with irradiated mouse embryonic fibroblast feeder cells | | | | | | | | | | | | |
| Media composition (mEpiSCs) | N2B27 media: 50% DMEM-F12, 50% Neurobasal, 0.5% N2 supplement, 1% B27 supplement without vitamin A, 2 mM glutamax, 1% penicillin-streptomycin, 0.1% 2-mercaptoethanol, 20 ng/ml activin A, 12.5 ng/ml heat stable bFGF and 175 nM NVP-TNKS656 | | | | | | | | | | | | |
| Passage at EpiLC-to-EpiSC conversion | P5-10 | P5-10 | P5-10 | P10 | P5 | P14 | P6 | P6 | P11 | P14 | P10 | P10 | P8 |
| Passage at experimental time | P5-10 | P5-10 | P5-10 | P17 | P13 | P20 | P12 | P12 | P18 | P21 | P17 | P17 | P15 |
| Mycoplasma results | Negative | Negative | Negative | Negative | Negative | Negative | Negative | Negative | Negative | Negative | Negative | Negative | Negative |
| Karyotyped by LC-WGS | | Yes - Normal | | | Yes - Normal | Yes - Normal | | | | | Yes - Normal | | Yes - Normal |

**Supplementary Table 4: Mouse embryonic stem cell lines**

| Genetic background | Biological replicates | Technical replicates |
| --- | --- | --- |
| B6129SF1/J | 3 | >10 |
| C57Bl/6J | 1 | 2 |
| C57Bl/6J X CAST/EiJ | 1 | 2 |

**Supplementary Table 5: EpiSC lines used in brain organoid experiments.**

### PRETHALAMUS

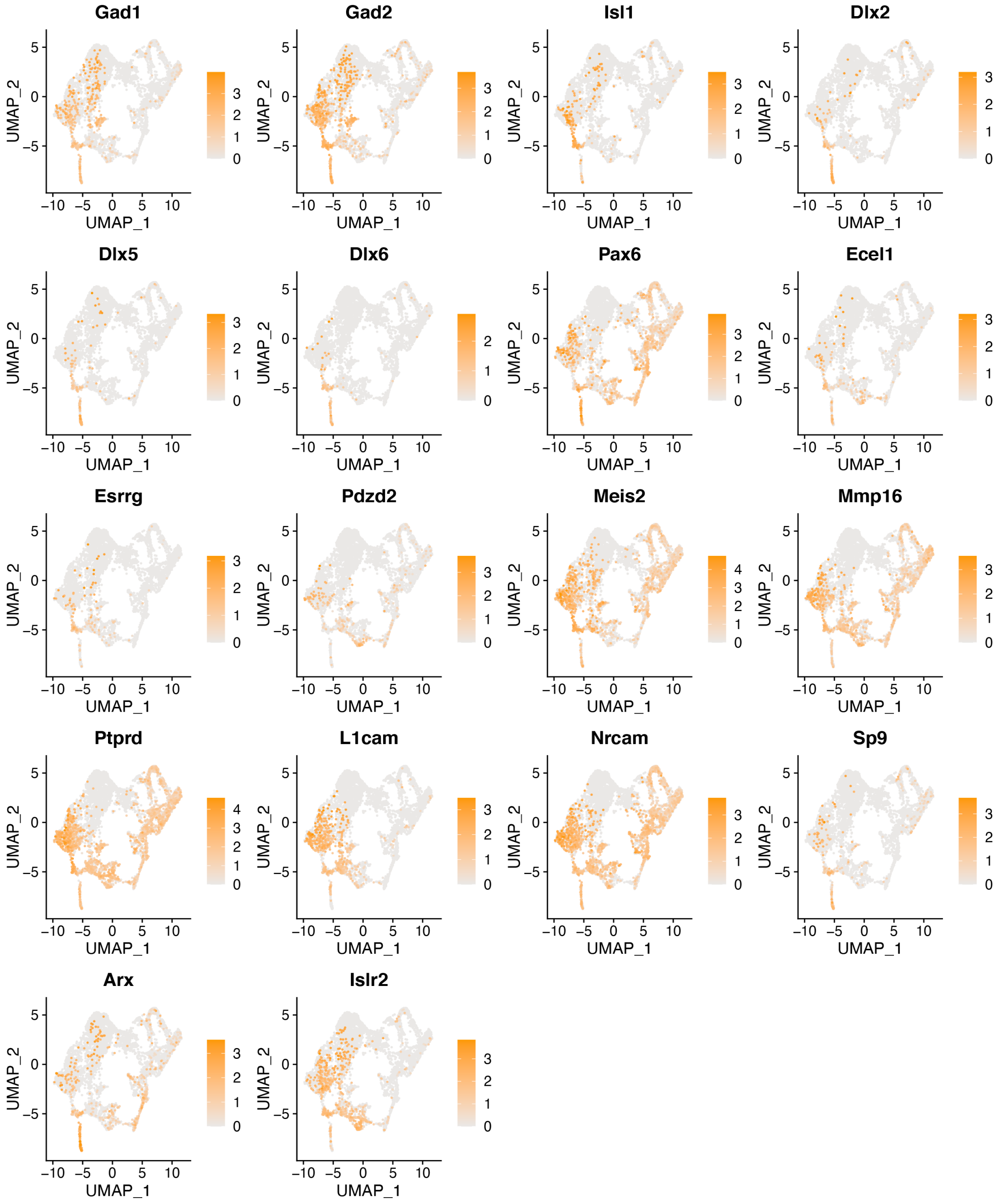

### CORTICAL HEM

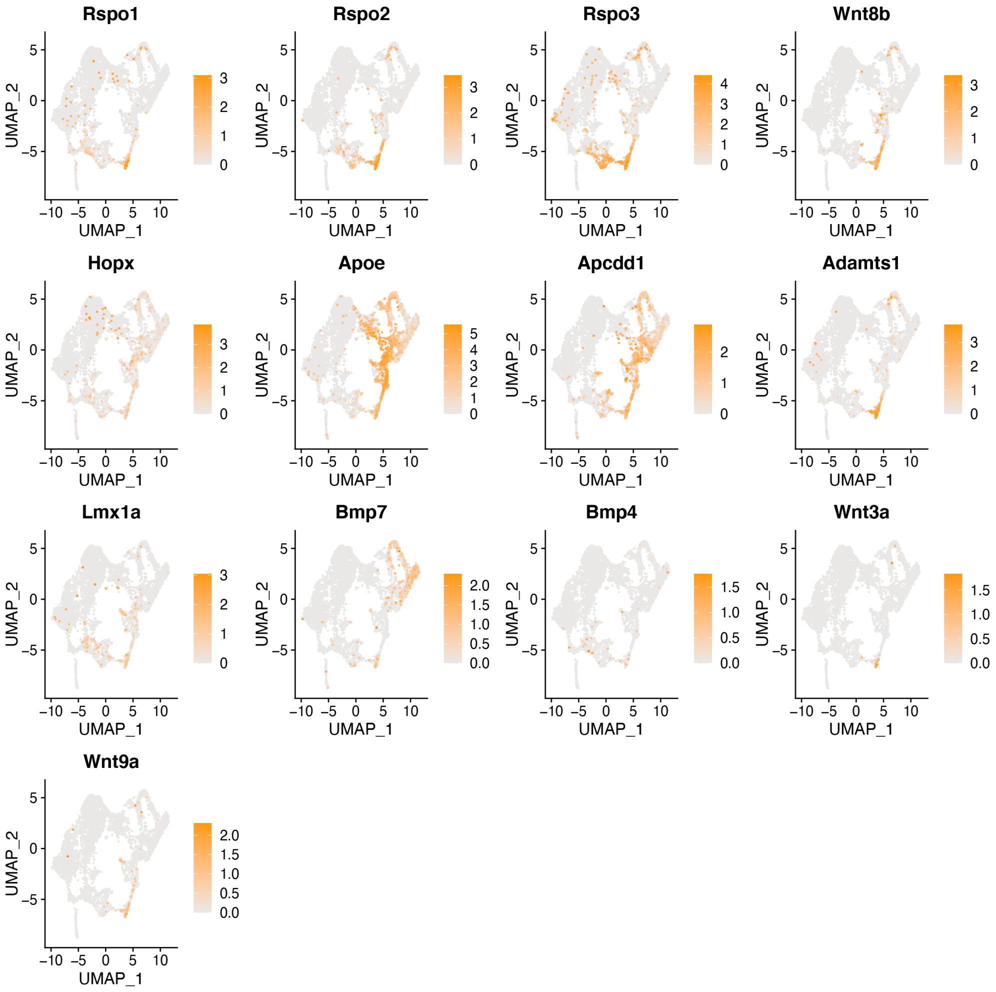

### HIPPOCAMPUS

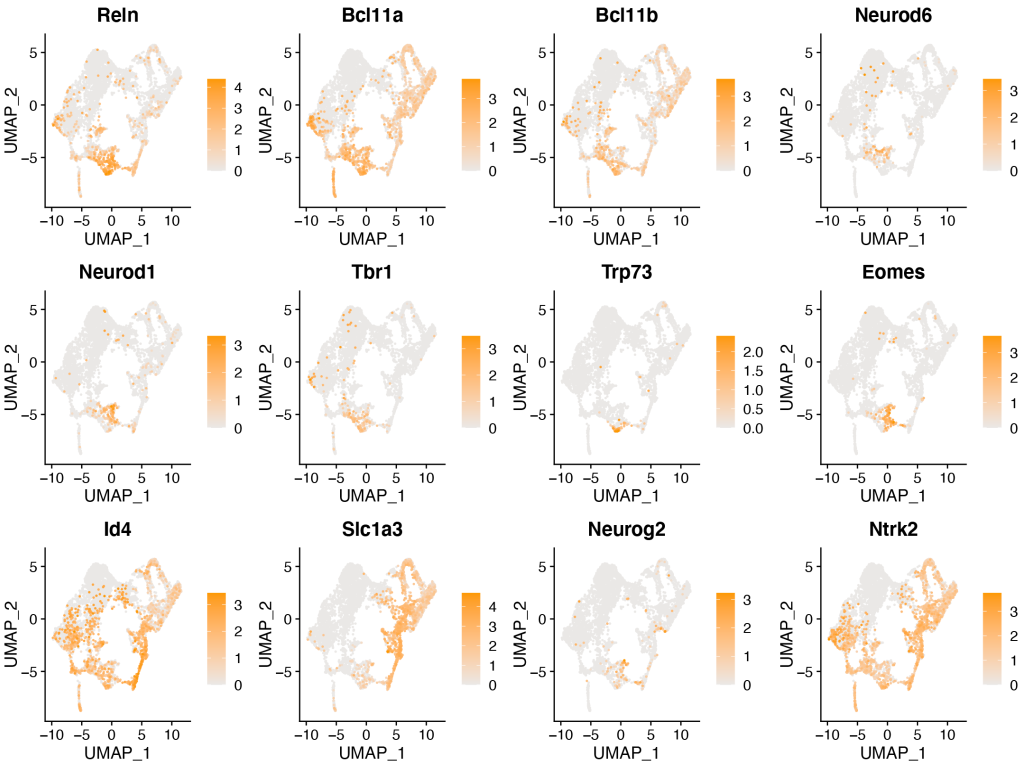


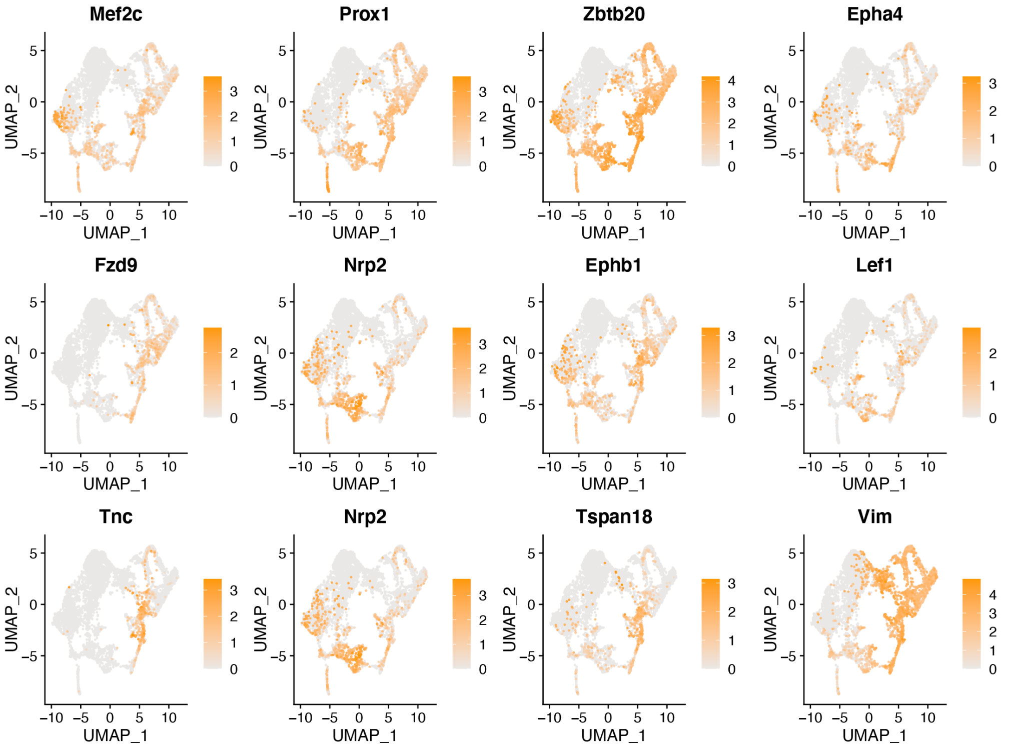

### OPCs

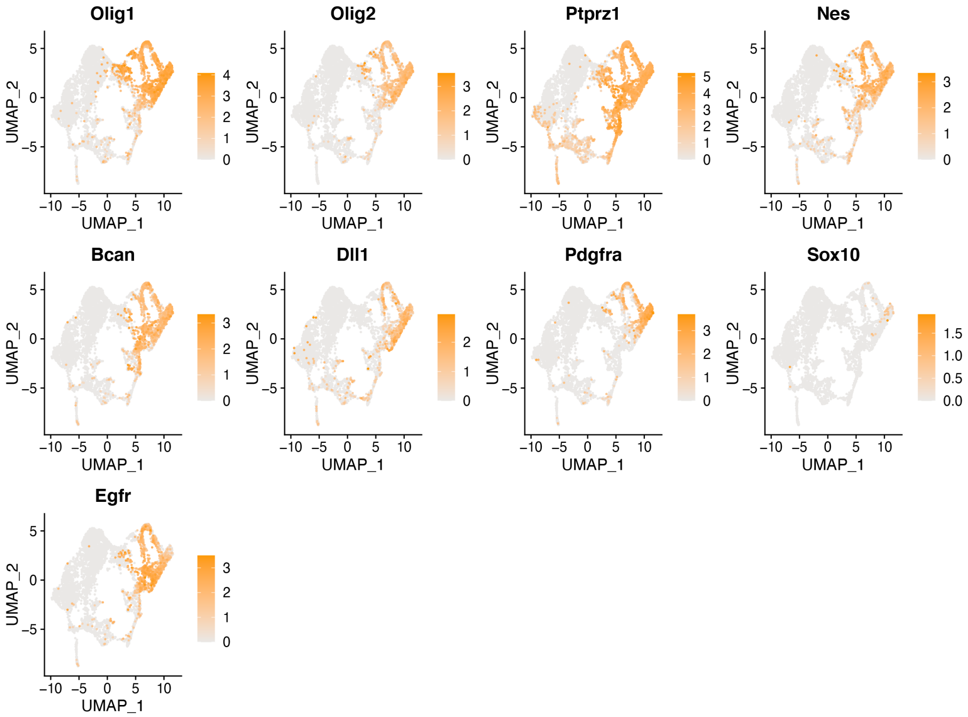


**Supplementary Table 6: Complete UMAPs, related to figure 6.**
