## Supplementary experimental methods for "Rapid and robust directed differentiation of mouse epiblast stem cells into definitive endoderm and forebrain organoids"

**Protocol: Mouse ESC to EpiSC conversion and EpiSC culture**

Notes before starting:

- Before use, all media should be warmed to 37 °C using a water or bead bath.
- All steps should be performed in a sterile BSC using rigorous aseptic technique.

Thawing ESCs

1. Remove cryovial of ESCs from liquid nitrogen storage and thaw rapidly with warmed media.
2. Once thawed, transfer into 15 mL falcon containing 5 mL Serum/Lif media (Table 2) to dilute the DMSO in the freezing media.
3. Centrifuge at 200 g for 3 minutes.
4. Aspirate media and resuspend cell pellet in appropriate volume of Serum/Lif media (Table 1).
5. Remove pre-plated feeders from the incubator and aspirate MEF media.
6. Add single cell suspension of ESCs to feeders and gently shake the plate left-to-right and up-and-down 3 times each to ensure cells are evenly distributed throughout.

Maintaining ESCs

*Important notes*

- Change media daily and passage cells upon 70% confluence at a 1:6 ratio.
- Feeders plated on gelatin should be prepared at least 8 hours in advance.

*Passaging ESCs*

1. Aspirate media and wash cells twice with PBS without calcium and magnesium ($PBS^{-/-}$).
2. Add warmed Accutase to wells for 2-3 minutes to dissociate the cells from the plate.
    a. *To speed up the dissociation process, place the plate with Accutase at 37 ℃ for 1-2 minutes.*
3. Gently tap the sides of the tissue culture plate to enhance cell dissociation.
4. Once all cells are lifted, collect Accutase and cells in a 15 mL centrifuge tube and dilute with equal amounts of $PBS^{-/-}$.
5. Centrifuge at 200 g for 3 minutes.
6. Aspirate supernatant and resuspend cell pellet in Serum/Lif media.
7. Remove MEF media from feeders and add the appropriate amount of ESC suspension to the plate for a 1:6 dilution.
8. If passaging long-term as ESCs, use collagenase to lift cells and accutase to bring to a single cell suspension to avoid accumulation of feeders. See "Passaging as EpiSCs" below.

ESC to EpiLC Conversion

*Prepare plate*

1. Coat the desired tissue culture plate with fibronectin diluted to 16.7 uL/mL in $PBS^{-/-}$. Rest plate at room temperature for ≥30 minutes.
2. Wash the coated plate twice with $PBS^{-/-}$.
3. Add additional $PBS^{-/-}$ to the plate until ready to add the cells to prevent desiccation. The plate can be stored with $PBS^{-/-}$ at 37 degrees for up to 4 hours.

*Create single cell suspension and plate*

1. Confirm ESC quality using a phase contrast microscope.
2. Aspirate ESC media from plate.
3. Use a serological pipette to wash each well twice with $PBS^{-/-}$.
4. Remove $PBS^{-/-}$ from the plate and apply pre-warmed Collagenase IV to each well to preferentially dissociate EpiSC colonies without feeder contamination.
    a. *Optional: place the plate with collagenase back in the incubator to increase the speed of dissociation.*
    b. *Gently tap the sides of the tissue culture plate to encourage colony lifting.*
    c. *Do not leave collagenase on wells for over 25-30 minutes as this will cause feeder cells to dissociate.*
5. Once the ESC colonies are lifted, gently collect colonies using serological pipette and transfer to a 15 mL centrifuge tube.
    a. *Optional: wash each well with HBSS++ to collect any remaining colonies and combine with the collagenase in the 15 mL tube.*
6. Centrifuge cells at 125 g for 4 minutes to pellet the ESC colonies and aspirate the supernatant.
7. Using a serological pipette, resuspend the pellet with 1 mL Accutase to break colonies into a single cell suspension. Leave the cell suspension under the hood for 2-4 minutes to ensure single cell suspension.
    a. *Optional: remove one drop of the suspension (~10 uL) and place on microscope slide. Observe with the microscope for residual colonies.*
8. Once the colonies are completely dissociated, add 5-10 mL $PBS^{-/-}$ to dilute the Accutase.
9. Centrifuge the solution at 200 g for 3 minutes to pellet single cells.
10. Aspirate supernatant and resuspend the pellet in EpiLC media (Table 2).
11. Count cells and plate at a density of 17,500 cells/$cm^2$.
12. Change the media 24 hours after plating. Cells should be cultured in EpiLC media for 48 hours before being converted to EpiSCs.

Conversion and Maintenance of EpiSCs

*Important notes*

- Change media daily and passage cells at 70% confluence, generally every other day
- Feeders plated on gelatin should be at least 8 hours before.

*Converting EpiLCs to EpiSCs*

1. After 48 hours of culture in EpiLC conditions, wash cells twice with $PBS^{-/-}$.
2. Add warmed Accutase to wells and collect in 15ml falcon tube once cells are lifted. Break the cells into small clumps of 5-10 cells (approx. 2-4 mins). Add equal volume of $PBS^{-/-}$ to dilute Accutase.
3. Centrifuge at 200 g for 3 minutes to pellet cells.
4. Resuspend in pre-warmed EpiSC media (Table 2).
5. Wash feeders twice with $PBS^{-/-}$. Aspirate the final PBS wash and plate 50,000-100,000 EpiLCs/$cm^2$ onto the feeders.
    a. *Note: Roughly, 1 well of a 12-well plate of EpiLCs can be seeded to 1 well of a 6-well plate for EpiSC culture.*

*Passaging EpiSCs*

1. Aspirate EpiSC media.
2. Use a serological pipette to wash each well twice with $PBS^{-/-}$.
3. Add pre-warmed Collagenase IV to each well to preferentially dissociate EpiSC colonies without feeder contamination.
    a. *Optional: place the plate with collagenase in the incubator to increase the speed of dissociation.*
    b. *Gently tap the sides of the tissue culture plate to encourage colony lifting.*
    c. *Do not leave collagenase on wells for over 25-30 minutes as this will cause feeder cells to dissociate.*
4. Once EpiSC colonies are lifted, gently collect colonies using serological pipette and transfer to a 15 mL centrifuge tube.
    a. *Optional: wash each well with HBSS++ to collect any remaining colonies and combine with collagenase in the 15 mL tube.*
5. Centrifuge cells at 125 g for 4 minutes to pellet EpiSC colonies and aspirate supernatant.
6. Using a serological pipette, resuspend the pellet with 1 mL Accutase to break colonies into clumps of 3-8 cells. Leave the cell suspension in hood for 1-2 minutes to ensure dissociation.
    a. *Optional: remove one drop of the suspension (~10uL) and place it on microscope slide. Observe colony size under the microscope.*
7. Once cells are dissociated, add 5-10 mL $PBS^{-/-}$ to dilute Accutase.
8. Centrifuge the solution at 200 g for 3 minutes to pellet cells.
9. Gently resuspend in EpiSC media to avoid additional dissociation.
10. Wash pre-plated feeders twice with $PBS^{-/-}$ to remove all MEF media and any traces of serum.
11. Plate EpiSCs on washed feeders with appropriate volume of media (Table 1) and place in incubator.
    a. *Optional: Check the size of the cell clumps after plating to ensure small colonies of 3-8 cells.*

12. EpiSCs grow very rapidly and therefore require daily media changes and are generally passaged every 48 hours.

Freezing EpiSCs

1. Aspirate media from each well of the plate and wash two times with $PBS^{-/-}$.
2. Add appropriate amount of accutase to each well (Table 1) and place in the incubator until the colonies are lifted.
3. Collect the mixture with a serological pipette and transfer to a 15 mL centrifuge tube.
4. Dilute Accutase with $PBS^{-/-}$.
5. Centrifuge cell solution at 200 g for 3 minutes.
6. Aspirate the supernatant.
7. Gently resuspend in EpiSC media with 10% DMSO, add to a cryotube and freeze.

Thawing EpiSCs

1. Remove EpiSCs from liquid nitrogen tank and thaw rapidly with warmed media.
2. Once thawed, add to a 15 mL falcon with 4 mL of N2B27 to dilute the DMSO.
3. Centrifuge at 200 g for 3 minutes to pellet cells.
4. Aspirate supernatant and gently resuspend the cells with EpiSC media.
5. Wash overnight feeders 2x with $PBS^{-/-}$ and plate cells directly onto feeders.

Frequent questions and concerns

Minimizing EpiSC differentiation (Figure 1B)

- Once there are many spontaneously differentiating colonies within the culture, it can be difficult to bring the line back to pluripotency. Prevention by culturing carefully is the best method to avoid spontaneous differentiation.
- Passage frequently, preferentially every other day, with no longer than 3 days on the same plate of feeders.
- Use collagenase to passage, as this will avoid propagation of differentiated cells.

**Protocol: Directed differentiation of mouse EpiSCs into definitive endoderm**

Notes before starting:

- Before use, all media should be warmed to 37 °C using a water or bead bath.
- All steps should be performed in a sterile BSC using rigorous aseptic technique.

Preparing the laminin coated cell culture dish

1. Dilute laminin-521 in PBS with calcium and magnesium to a final concentration of 10 ug/mL. Optionally, laminin from EHS murine sarcoma basement membrane (Sigma, L2020) can be used as a replacement at 20 ug/mL.

2. Once combined, use a serological pipette to dispense the diluted laminin into each well of a tissue culture plate (see Table 3 for volume recommendations). Shake plate to ensure that the entire surface is covered.
3. Place the laminin-coated tissue culture plate(s) in cell culture incubator at 37°C for 2 hours or O/N at 4°C.

<u>Preparing and plating EpiSCs</u>

1. Confirm EpiSC quality using a phase contrast microscope (Figure 1A).
2. Aspirate EpiSC media from plate.
3. Use a serological pipette to wash each well twice with $PBS^{-/-}$.
4. Remove the $PBS^{-/-}$ from the plate and apply pre-warmed Collagenase IV to each well to preferentially dissociate EpiSC colonies without feeder contamination.
    a. *Optional: place the plate with collagenase back in the incubator to increase the speed of dissociation.*
    b. *Gently tap the sides of the tissue culture plate to encourage colony lifting.*
    c. *Do not leave collagenase on wells for over 25-30 minutes as this will cause feeder cells to dissociate as well and should be avoided.*
5. Once the EpiSC colonies are lifted, gently collect colonies using serological pipette and transfer to a 15 ml centrifuge tube.
    a. *Optional: wash each gently with HBSS++ to collect any remaining colonies and combine with collagenase in the 15 mL tube.*
6. Centrifuge cells at 125 g for 4 minutes to pellet EpiSC colonies and aspirate the supernatant.
7. Using a serological pipette, resuspend the pellet with 1 mL Accutase to break colonies into a single cell suspension. Leave the cell suspension in hood for 2-4 minutes to ensure single cell suspension
    a. *Optional: remove one drop of the suspension (~10 uL) and place on microscope slide. Observe under the microscope for residual colonies.*
8. Once cells are completely dissociated, add 5-10 mL $PBS^{-/-}$ to dilute Accutase.
9. Centrifuge the solution at 200 g for 3 minutes to pellet single cells.
10. Aspirate the supernatant, resuspend in plating media (Table 4) and count the number of cells.
11. Calculate the appropriate volume of media and number of cells to seed the desired plate, then prepare master solution with 110,000 cells/$cm^2$ (Table 3).
12. Remove the laminin coated plate from incubator and aspirate laminin solution. Do not allow plate to dry—the cell suspension must be added immediately.
13. Homogenize the cell solution and immediately add the cell suspension to the plate.
14. Place the plate in the incubator for 5-6 hours. During this time cells will attach to the plate.

Changing media

1. After 5-6 hours, gently remove the plate from the incubator and check under microscope to confirm that most cells have attached. At this stage, the cells can be easily washed off if media is added too vigorously.
2. Slowly aspirate media off all wells at a 45-degree angle to the BSC surface. Gently add $PBS^{-/-}$ to wash off any residual media.
    a. *IMPORTANT: Use one hand to tilt the tissue culture plate towards you at a 45° angle and maintain this angle while gently removing and dispensing media into the side of the well (slow ejection speed on the pipettor). This will ensure minimal loss of cells.*
3. Using same technique, add appropriate amount of Differentiation Media 1 (Table 4).
4. Place the tissue culture dish in the incubator and allow to sit for 16 more hours.
5. After 16 hours, aspirate Differentiation Media 1.
6. While resting serological pipette tip on bottom wall of well, gently add $PBS^{-/-}$ to dilute any residual media.
7. Using same technique as $PBS^{-/-}$ application, add appropriate amount of Differentiation Media 2 (Table 4).
8. Return the tissue culture dish to the incubator for 24 hours.
9. After 24 hours in Differentiation Media 2, the cells should have reached the definitive endoderm stage. Examine all wells to confirm appropriate density and morphology.

Fixing and immunostaining

1. To fix cells, wash each well two times with $PBS^{-/-}$.
2. Add 4% PFA to each well and allow to rest at room temperature for 15 minutes.
3. After 15 minutes, aspirate the PFA and wash twice with $PBS^{-/-}$.
4. Add $PBS^{-/-}$ to each well and refrigerate at 4 °C until staining.

Frequent questions and concerns

Low cell density after plating

- Ensure the laminin was diluted and left for specified amount of time (2 hours). Do not rinse culture plates after laminin coating.
- Some cell lines have weak attachment. If this is a recurring issue for specific lines, increase the starting density when plating.

Cell loss during media changes

- Use one hand to tilt the tissue culture plate towards you at a 45° angle and maintain this angle while gently removing and dispensing media into the side of the well (slow ejection speed on the pipettor). This will ensure minimal loss of cells.
- Increase the size of the wells you are using. Cell loss is more common on smaller sized wells.
- Wait a longer period of time (6-8 hours) before changing plating media.

Low efficiency of differentiation

- It is important to begin with high quality EpiSCs with minimal spontaneous differentiation. At the beginning of the directed differentiation protocol, EpiSCs colonies should number ~50-200 and the plate should be no more than 70-80% confluent.
- This protocol was optimized on EpiSCs passaged no more than 18 passages as EpiSCs and it is unknown how high passage lines may respond to the differentiation protocol. In general, it is best to begin with low-passage ESCs to convert to EpiSCs and avoid prolonged culture.

Timing

- The timing of each media change was optimized to produce the highest Sox17 and FoxA2 expression at 40 hours of culture. Changing the timing of Step 1 and Step 2 can lead to reduced efficiency.

**Protocol: Generation of forebrain organoids from mouse EpiSCs**

<u>Notes before starting:</u>

- Before use, all media should be warmed to 37 °C using a water or bead bath.
- All steps should be performed in a sterile BSC using rigorous aseptic technique.

<u>Formation of EBs</u>

1. Confirm EpiSC quality using a phase contrast microscope (Figure 1A).
2. Aspirate EpiSC media from plate.
3. Use a serological pipette to wash each well twice with $PBS^{-/-}$.
4. Remove the $PBS^{-/-}$ from the plate and apply pre-warmed Collagenase IV to each well to preferentially dissociate EpiSC colonies without feeder contamination.

- *Optional: place the plate with collagenase back in the incubator to increase the speed of dissociation.*
- *Gently tap the sides of the tissue culture plate to encourage colony lifting.*
- *Do not leave collagenase on wells for over 25-30 minutes as this will cause feeder cells to dissociate as well and should be avoided.*

5. Once the EpiSC colonies are lifted, gently collect colonies using serological pipette and transfer to a 15 mL centrifuge tube.

- *Optional: wash each gently with HBSS++ to collect any remaining colonies and combine with collagenase in the 15 mL tube.*

6. Centrifuge cells at 125 g for 4 minutes to pellet EpiSC colonies and aspirate the supernatant.
7. Using a serological pipette, resuspend the pellet with 1 mL Accutase to break colonies into a single cell suspension. Leave the cell suspension in the hood for 2-4 minutes to ensure single cell suspension.

- *Optional: remove one drop of the suspension (~10 uL) and place on microscope slide. Observe under the microscope for residual colonies.*

8. Once cells are completely dissociated, add 5-10 mL $PBS^{-/-}$ to dilute Accutase.
9. Centrifuge the solution at 200 g for 3 minutes to pellet single cells.
10. Aspirate the supernatant, resuspend in EB formation media (Table 5) and count cells.
11. Calculate the appropriate volume of media and number of cells to seed the desired number of wells, then prepare master solution with 1,200,000 cells/well of Aggrewell, with each well containing a total of 2 mL media.
12. Using a serological pipette, disperse cell solution into well(s) of Aggrewell.
13. Using a P1000 pipette, gently pipet up and down to equally distribute the cells within the well(s).
14. Centrifuge the Aggrewell plate at 100 g for 3 minutes. Confirm cells are evenly distributed using microscope.
15. Place Aggrewell in incubator for 24 hours to form EBs. Avoid moving plate during the EB formation phase.

<u>Collection and embedding of organoids</u>

*Important note:*

- Matrigel should be thawed overnight on ice at 4°C (or for at least 3-4h)

1. Following 24 hours in EB formation media, Aggrewell should contain uniform, well-formed developing organoids. Using a P1000 pipette, gently remove as much media as possible from the Aggrewell without disturbing organoids.
2. Using a P1000, add 1 mL of Neural Induction media (Table 5) to Aggrewell and, using a wide orifice pipette tip, gently resuspend the developing organoids.
3. Transfer organoids to a 6 cm Petri dish containing 5 mL Neural Induction media and swirl briefly to dilute EB formation media. Transfer organoids to a 10 cm Petri dish containing 10 mL Neural induction media.
   a. *Note: washing with $PBS^{-/-}$ should be avoided as this may cause organoids to adhere to the plate.*
4. Determine the total number of organoids desired and how many wells of a low-adherence 6 well plate are needed.
   a. *Note: Each well of the plate should contain 20-50 organoids.*
5. Then, carefully gather 20-50 organoids in 67 uL of Neural Induction media and mix with 100 uL of Matrigel. Carefully dispense the mix of Matrigel, media, and organoids in the center of a well, being sure to avoid the walls of the well (Derived from Qian et al., 2018).
6. Repeat for desired number of wells then place the plate in the incubator for 30 minutes to solidify the Matrigel.
7. Following the 30-minute incubation, carefully remove the plate from the incubator and gently add 3 mL of Neural Induction media to each well. Return to incubator for 24 hours.
   a. *Note: when adding the media, Matrigel domes may float off plate. This does not affect organoid quality.*

Maintenance and patterning of organoids

1. Following 24 hours in Neural Induction media, gently remove as much media as possible from the well without disturbing Matrigel domes.
2. Add 3 mL of Neuroepithelial Expansion media (Table 5) to each well. Place plate in incubator for 48 hours.
3. After 48 hours in Neuroepithelial Expansion media, gently remove media and wash twice with $PBS^{-/-}$.
    a. *Note: Wash and media change should be performed under a dissection microscope to avoid organoid loss.*
4. After removing second $PBS^{-/-}$ wash, add 2mL Corning Cell Recovery Solution to each well of the 6 well plate. Gently swirl plate and place in incubator for 30 minutes to remove Matrigel from organoids.
    a. *Optional: To improve recovery, gently swirl plate every 5-10 min while in incubator.*
5. Following Matrigel removal, gently add 2 mL of Neuronal media (Table 5) to each well of the 6 well plate to dilute solution.
6. Using a wide orifice pipette tip, transfer the organoids to a freshly prepared low adherence 6 well plate containing 2 mL Neuronal media per well.
7. Using a wide orifice pipette tip, transfer the organoids to an additional freshly prepared low adherence 6 well plate containing 2 mL Neuronal media per well to ensure maximum dilution of Corning Cell Recovery solution.
8. Finally, using a wide orifice pipette tip, transfer organoids to a 10 cm Petri dish containing 12.5mL of Neuronal media and place on shaker.
    a. *Note: speed of shake will depend on the specific shaker being used. For Celltron benchtop shaker, 65 rpm was sufficient in preventing organoid merging while minimizing impact.*
9. Change media using fresh Neuronal media every other day until collection day.

**Protocol: Immunostaining of mouse EpiSC-derived forebrain organoids (method derived from Dekkers et al., 2019)**

Fixation and immunostaining of organoids

1. Coat a 15 mL falcon tube with 1% BSA to avoid organoid attachment and place on ice.
2. While on ice, gently add organoids to the pre-coated falcon tube and wash three times with 4°C $PBS^{-/-}$.
    a. *Note: Organoids at day 4 or earlier should be centrifuged at 70 g for 3 minutes at 4 ℃ for each wash step. Organoids later than day 4 will naturally settle to the bottom of the tube due to their size.*
3. To fix organoids, add 4% PFA to 15 mL falcon tube until organoids are completely submerged. Place on ice for 45 minutes and gently resuspend halfway through incubation to ensure proper fixation.
4. Following fixation, wash three times with $PBS^{-/-}$ containing 0.1% Triton-X (PBST) to remove any remaining PFA.

5. To permeabilize organoids, add $PBS^{-/-}$ containing 0.5% Triton-X until organoids are completely submerged and incubate for 15 minutes at 4°C.
6. Following 15-minute incubation, remove permeabilization buffer and block organoids for 15 minutes in organoid washing buffer (OWB) containing 0.2% Triton-X, 0.02% SDS and 0.2% BSA in $PBS^{-/-}$. Calculate appropriate volume of OWB by determining number of staining conditions (200 uL OWB per staining condition).
7. Equally distribute the organoids in a low-adhesion 24-well plate with a total of 200 uL OWB and desired number of organoids in each well. Each well will represent a single staining condition.
8. Following 15-minute blocking, directly add primary antibodies of interest prediluted in 200 uL OWB to each well for a total volume of 400 uL per well.
9. Place plate on shaker and incubate in primary antibodies overnight at 4°C.
10. Following overnight incubation, add 800 uL of OWB directly to each well and place on shaker at room temperature for 5 minutes.
11. Next, remove 1 mL of solution, replace with a fresh 1mL of OWB, and place on shaker for two hours at room temperature to wash organoids.
12. Repeat this wash step two additional times, for a total of 6 hours of washing.
13. After 6 hours of washing, remove 1 mL of OWB from each well and directly add 200 uL of prediluted secondary antibodies and DAPI (final concentration of 1 μM) in OWB, for a total volume of 400 uL per well.
14. Place on shaker and incubate overnight at 4°C.
15. Following overnight secondary antibody incubation, remove from 4°C, and add 800 uL of OWB with 1 μM DAPI per well. Place on shaker at room temperature and allow to shake for 5 minutes.
16. Next, remove 1 mL of OWB from each well and replace with a fresh 1 mL of OWB with 1 μM DAPI. Place on shaker at room temperature for 2 hours to wash organoids.
17. Repeat this wash step twice more, for a total of 6 hours of washing.

<u>Preparation of slide for imaging</u>

1. Using a standard hole puncher, punch a hole (0.5cm Ø) into a sticky silicone pad (20x20x1 mm).
2. Place sticky pad on imaging slide. (Figure 2)

<u>Mounting of organoids for imaging</u>

1. Using a wide orifice pipette tip, Transfer the organoids to a 1.5 mL Eppendorf.
2. Remove as much supernatant as possible without disturbing the organoids
    a. *Note: If the organoids at day 4 or less, centrifuge at 70 g for 3 minutes to pellet.*
3. Using a wide orifice pipette tip, add 35 uL of DeepClear solution, gently resuspend organoids, and transfer to prepared slide. Add a coverslip and image.

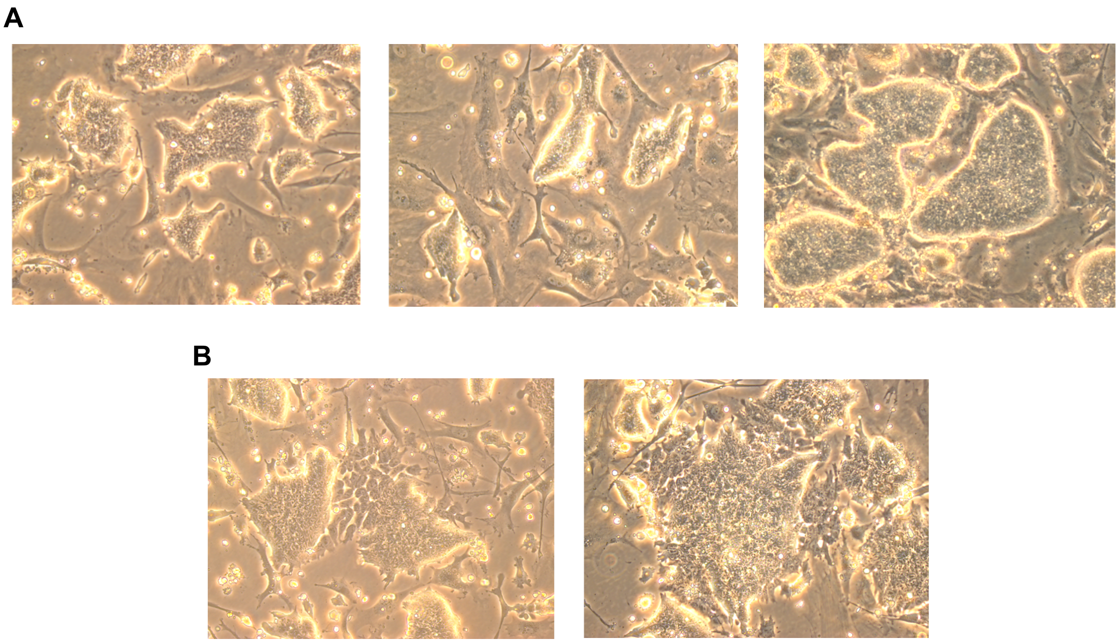


**Figure 1: EpiSCs in culture.** (A) Standard culture of EpiSCs. (B) Spontaneous differentiation in EpiSC culture.

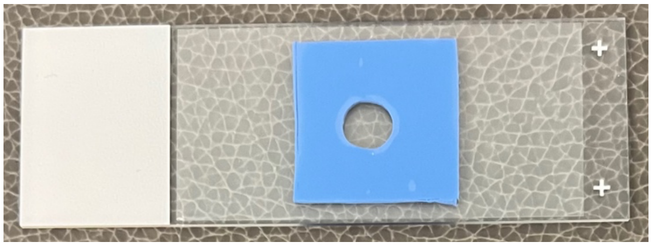

**Figure 2: Slide for organoid imaging.**

**Table 1.** Plate sizes and suggested volumes for stem cell culture

| Plate size | Surface area ($cm^2$) | Volume of collagenase, $PBS^{-/-}$ and Accutase (mL) | Media Volume (mL) |
|---|---|---|---|
| 96 well | 0.32 | 0.1 | 0.2 |
| 48 well | 1.1 | 0.3 | 0.6 |
| 24 well | 1.9 | 0.4 | 0.8 |
| 12 well | 3.5 | 0.5 | 2 |
| 6 well | 9.6 | 1 | 5 |
| 10cm dish | 56.7 | 5 | 15 |

**Table 2.** Media composition for stem cell culture

| MEF | ESC | EpiLC | EpiSC |
|---|---|---|---|
| MEF media | S/L media | N2B27 | N2B27 |
| | 3uM CHIR99021* | 12.5 ng/ml Heat stable recombinant human bFGF | 12.5 ng/ml Heat stable recombinant human bFGF |
| | 1uM PD0325901* | 20 ng/ml Activin A | 20 ng/ml Activin A |
| | | 1% KOSR | 175 nM NVP-TNKS656 |

*Optional components to create 2i/SL culture conditions

**Table 3.** Plate sizes and suggested volumes for DE differentiation

| Plate size | Surface area ($cm^2$) | Volume of laminin, collagenase, PBS (ml) | Media Volume (ml) | Number of cells per well |
|---|---|---|---|---|
| 96 well | 0.32 | 0.1 | 0.2 | 35,000 |
| 48 well | 1.1 | 0.3 | 0.6 | 120,000 |
| 24 well | 1.9 | 0.4 | 0.8 | 208,000 |
| 12 well | 3.5 | 0.5 | 2 | 382,000 |
| 6 well | 9.6 | 1 | 5 | 1,050,000 |
| 10cm dish | 56.7 | 5 | 15 | 6,200,000 |

**Table 4.** DE differentiation media composition

| Plating Media | Differentiation media 1 | Differentiation media 2 |
|---|---|---|
| CDM + 0.7 μg/ml insulin | CDM + 0.7 μg/ml insulin | CDM + 0.7 μg/ml insulin |
| 12.5 ng/ml FGF | 3 μM CHIR99021 | 100 ng/ml Activin A |
| 20 ng/ml Activin | 40 ng/ml Activin A | 100 nM LDN-193189 (BMP inhibitor) |
| 175 nM NVP-TNKS656 | | |
| 1% Knockout Serum Replacement | | |
| 2 μM Thiazovivin | | |

**Table 5.** Media composition for neural organoids

| EB Formation media | Neural induction media | Neuroepithelial expansion media | Neuronal media |
|---|---|---|---|
| N2B27 (B27 without vitamin A) | N2B27 (B27 without vitamin A) | N2B27 (B27 without vitamin A) | N2B27 (B27 with vitamin A) |
| 50 nM chroman-1 | 100 nM LDN | 100ng/ml Fgf8b | 20ng/mL BDNF |
| 5 μM emricasan | 10 μM SB431542 | | 20ng/mL GDNF |
| 100 nM LDN | 100 nM PD173074 | | |
| 10 μM SB431542 | 4 nM LGK974 | | |
| 100 nM PD173074 | | | |
| 4 nM LGK974 | | | |

**MEF MEDIA**: [**For 500ml**] Sterile filter and store at 4°C

| Solution (stock concentration) | Volume | Final concentration |
|---|---|---|
| DMEM w/ glutamine (1x) | 445 mL | 2 mM (L-glutamine) |
| Penicillin-streptomycin (100x) | 5 mL | 100 ug/mL |
| FCS (Hyclone) | 50 mL | 10% |

**SERUM/LIF MEDIA**: [**For 500ml**] Sterile filter and store at 4°C

| Solution (stock concentration) | Volume | Final concentration |
|---|---|---|
| DMEM w/ glutamine (1x) | 430ml | 2mM (L-glutamine) |
| NEAA (100x) | 5ml | 0.1mM |
| Sodium pyruvate (100x) | 5ml | 1mM |
| Penicillin-streptomycin (100x) | 5ml | 100ug/ml |
| 2-mercaptoethanol (1000x) | 500ul | 0.1mM |
| FCS (Hyclone) | 50ml | 10% |
| ESGRO LIF (100x) | 5ml | 1000U/ml |

**N2B27 MEDIA**: [**For 500ml**] Sterile filter and store at 4°C

| Solution (stock concentration) | Volume | Final concentration |
|---|---|---|
| DMEM- F12 | 241ml | |
| Neurobasal medium | 241ml | |
| N2 supplement | 2.5ml | 1:200 |
| B27 supplement | 5ml | 1:100 |
| GlutaMax | 1.875ml | 2mM |
| Penicillin-streptomycin (100x) | 5ml | 100ug/ml |
| 2-mercaptoethanol (1000x) | 500ul | 0.1mM |

**CHEMICALLY DEFINED MEDIA (CDM)**: [**For 500ml**] Sterile filter and store at 4°C

| Solution (stock concentration) | Volume | Final Concentration |
|---|---|---|
| IMDM (1x) | 243 ml | 50% |
| F12 with GlutaMAX (1x) | 243 ml | 50% |
| Chemically Defined Lipid Concentrate (100x) | 5 ml | 1x |
| Monothioglycerol (11.5 M) | 19.3 μL | 450 μM |
| Polyvinyl Alcohol (100 mg/mL) | 5 ml | 1 mg/mL |
| Apo transferrin (10 mg/mL) | 750 μL | 15 ug/mL |
| Glutamax (100x) | 2.5 ml | 100x |
| Insulin (10 mg/mL) | 35 μL | 0.7 μg/ml |

### Preparation of stock solutions

**Activin A: Peprotech (120-14P)**

- Reconstitute lyophilized protein to 100ug/ml in sterile $H_2O$ with 0.1% BSA (5,000X solution)
- Aliquot and store at -80 C for up to 3 months.

**Heat-stable bFGF2: Gibco (PHG0360)**

- Reconstitute FGF to 1 g/L in sterile diH2O.
- Add sterile PBS + 0.1% BSA to dilute solution to 25 ug/ml (2,000X stock for a working concentration of 12.5 ng/mL).
- Store at -20 for up to 12 months.

**NVP-TNKS656: Selleck chemicals (S7238)**

- Dilute powder with DMSO to make a 35 mM solution (for 10 mg, add 577 μL DMSO)
- Dilute 35 mM stock further to 350 uM
- Aliquot 350uM stock (2,000X) and store at -80 C for up to 2 years.

**Collagenase Type IV: Gibco (17104109)**

- Add 10 ml of HBSS++ directly to the vial of collagenase.
- Vortex to complete dissolution.
- Determine final volume of HBSS++ required to bring collagenase solution to 5 u/μL (10X) & vortex.

- Filter to sterilize stock solution with a low protein binding filtration unit (PES Membrane, 0.2 μm).
- Aliquot and store at -20C for up to 24 months.
- Add HBSS++ to bring concentration to 5X before use.

**Thiazovivin: Sigma (SML1045)**
- Resuspend in DMSO to make 2mM stock (for 5 mg, 8.030 mL DMSO for a 1,000X solution).
- Aliquot and store at -20C for up to 6 months.

**CHIR99021: Tocris Bioscience (252917-06-9)**
- Resuspend in DMSO to make 5mM stock (for 10mg, 4.2979 mL DMSO for a 5,000X working solution).
- Aliquot and store at -20C.

**Apo-transferrin: InVitra (777TRF029)**
- Resuspend 1g in 100 mL diH2O, mix until complete homogenization and filter.
- This will give a 666 X stock solution. Aliquot and store at -80C.

**Lif: Sigma (ESG1107)**
- Each vial contains $10^7$ units/mL. Dilute it 100 times in sterile tissue culture media to make a 100x solution, which is stored at 4C until use.

**Insulin: Sigma (91077C)**
- Take 250 mg, add 25 mL diH2O, leading to a cloudy solution.
- Add HCl until a pH=3.0 is achieved.
- Filter and store at 4C.

**LDN-193189: Stemgent (04-0074)**
- Dilute the powder (2 mg) with 22.75 mL DMSO to make a 200 μM solution (2,000X).
- Aliquot and store at -20C for up to 6 months.

**PD0325901: Tocris (4192)**
- Add 830 μL of DMSO to 10mg powder to make a 25 mM stock solution.
- Aliquot and store at -20C.

**Polyvinyl alcohol (PVA): Sigma (341584)**
- Within a clean, autoclaved bottle (around 2 L capacity), add 20 g of PVA and 200 mL diH2O (100 mg/mL stock, 100X).
- Autoclave (wet settings) with the cap slightly opened for 1-2h until the powder fully dissolves.
- Store at 4C and use it within the next 3 months.

**Chroman 1: MedChem Express (HY-15392)**
- Dilute the powder (5 mg) with 11.4548 mL DMSO to make a 1 mM solution (20,000X).
- Aliquot and store at -80C for up to 6 months.

**Emricasan: Selleckchem (S7775)**
- Dilute the powder (25 mg) with 877 µL of DMSO to make a 50mM solution (10,000X).
- Aliquot and store at -80C for up to 2 years.

**SB-431542: Tocris (1614)**
- Dilute the powder (10 mg) with 2.6 mL 100% ethanol to make a 200 µM solution (1,000X).
- Aliquot and store at -20C for up to 1 month.

**PD-173074: STEMCELL technologies (72164)**
- Dilute the powder (10 mg) with 19.09 mL of DMSO to make a 1 mM solution (10,000X).
- Aliquot and store at -80C for up to 2 years.

**LGK-974: Selleck Chemicals (S7143)**
- Dilute the powder (1 mg) with 63.06 mL DMSO, warmed at 50C in a water bath if needed to fully dissolve, to make a 40 µM solution (10.000X) .
- Aliquot and store at -80C for up to 2 years.

**FGF8b: R&D systems (423-F8-025/CF)**
- Dilute the powder (25 µg) in 250 uL PBS-/- to make a 1,000X solution, store at 4C for up to 3 months.

**BDNF: Peprotech (450-02)**
- Resuspend the powder (50 µg) in 2.5 mL PBS-/- with 0.1% BSA to make a 1.000X solution.
- Aliquot and store at -20C for up to a year.

**GDNF: Peprotech (450-10)**
- Resuspend the powder (50 µg) in 2.5 mL PBS-/- with 0.1% BSA to make a 1.000X solution.
- Aliquot and store at -20C for up to a year.
